# Neutrophils are critical for placental and fetal infection with the human pathogen *Listeria monocytogenes*

**DOI:** 10.64898/2026.03.16.712023

**Authors:** Myriam Ripphahn, Nikita Raj, Hellen Ishikawa-Ankerhold, Zhouyi Rong, Bastian Popper, Bianca Portugal Tavares de Moraes, Xia Li, Ying Chen, Roland Immler, Lou-Martha Wackerbarth, Dominic van den Heuvel, Barbara Schmidinger, Rainer Haas, Udo Jeschke, Sven Kehl, Anne Loesslein, Steffen Massberg, Farida Hellal, Ali Ertürk, Markus Sperandio

**Author notes:** Equal contribution (Myriam Ripphahn, Nikita Raj). Correspondence: Markus Sperandio, M.D., Institute of Cardiovascular Physiology and Pathophysiology, Biomedizinisches Centrum München, Ludwig-Maximilians-Universität, Großhaderner Str. 9, 82152 Planegg-Martinsried, GERMANY, voice.

## Abstract

Fetal listeriosis, caused by *Listeria monocytogenes* (Lm), represents a severe infectious disease. During pregnancy, the unique intrauterine environment permits Lm to breach the maternal-fetal barrier by infiltrating the placental trophoblast layer and ultimately leading to significant fetal morbidity and mortality. However, the exact pathways remain incompletely understood. In this study, we sought to elucidate the molecular mechanisms underlying the early stages of placental and fetal Lm infection. Our results reveal that neutrophils serve as a survival niche for Lm within the vasculature, facilitating placental and fetal infection in the mouse *in vivo*. This finding was further substantiated by *in vitro* studies using TLR2-stimulated trophoblast cells and Lm-infected neutrophils. Additionally, tissue-clearing analysis of Lm-infected placentas demonstrated neutrophil colocalization with Lm is rare within placental tissue. Collectively, our findings identify maternal neutrophils as an essential viability niche for Lm enabling their entry into trophoblast cells, thereby promoting placental and fetal infection.

**One Sentence Summary:** Maternal circulating neutrophils act as survival niche for *Listeria monocytogenes* and facilitate placental/fetal infection with *Listeria* during pregnancy.

## Introduction

The facultative intracellular Gram-positive bacterium *Listeria monocytogenes* (Lm) is a foodborne ubiquitously distributed pathogen and the causative agent for listeriosis (Bucur et al., 2018; Koopmans et al., 2023). Listeriosis is an infectious disease occurring worldwide underlined by current cases of disease outbreaks in North America, South Africa, Australia, and Europe (Alland et al., 2025; Disson et al., 2025; Fernandez-Martinez et al., 2022; Lachmann et al., 2022; Pereira et al., 2023; Samut et al., 2025; Thomas et al., 2020). Discovered in 1926 by Murray as a rabbit pathogen, decades of research have established Lm as a model organism to study host-pathogen interactions. Lm is able to overcome many barriers, including the intestinal, blood-brain, and placental barrier (Charlier et al., 2020). But, despite intense research, the initial mechanism by which Lm overcomes the maternal-fetal barrier and subsequently infects the fetus still remains unclear. The consequences can be severe and pose a relevant health problem globally, as Lm infection is particularly risky during pregnancy (Kraus et al., 2024), and can lead to preterm labor and birth, fetal and neonatal sepsis with high morbidity and mortality (Charlier et al., 2022; LaTuga, 2025; Villa et al., 2025).

During pregnancy, the immunological environment in the mother and fetus is unique and maternal immune cell interaction with the placenta needs to be strictly organized to avoid rejection of the semi-allogenic fetus on one hand and to provide protection to the mother and fetus against invading pathogens on the other hand (Ander et al., 2019; Moffett and Loke, 2006). To optimally adapt to the specific needs of the fetus during pregnancy, the composition and function of maternal immune cell trafficking in and out of the placenta are tightly controlled and a prerequisite for successful pregnancy (Ander et al., 2019; Giaglis et al., 2016; PrabhuDas et al., 2015).

The maternal-fetal barrier in the placenta is the critical interface between the maternal and fetal tissues, responsible for regulatory and immune barrier functions that adapt across gestation (O’Brien and Wang, 2023). This barrier consists of the uterine wall-associated decidua on the maternal side and a special cell type, called trophoblasts on the fetal side (Knofler et al., 2019). Trophoblasts originate from the trophoectoderm of the blastocyst (Ander et al., 2019). In the human hemochorial placenta, as well as in the labyrinthine placenta of mice, several types of trophoblast cells are present (Ander et al., 2019). Syncytiotrophoblast cells (SYN) are in direct contact with maternal blood and are primarily responsible for gas and nutrient exchange (PrabhuDas et al., 2015). SYN form a multinucleated syncytial layer above the mononucleated cytotrophoblast (CTB) layer and a basement membrane underneath (Knöfler *et al*., 2019). CTBs can differentiate into SYNs or into another mononucleated trophoblast cell type, extravillous cytotrophoblasts (EVT), which is critical for anchoring the villous tree to the decidua at the start of the second trimester (Ander et al., 2019; Turco and Moffett, 2019). EVTs remodel the lumen of spiral arteries and interact with decidual leukocytes, supporting placental development, but also rendering the organ more susceptible to pathogens such as Lm (Semmes and Coyne, 2022; Silva and Serakides, 2016).

Several mechanisms have been proposed on how Lm survives in the maternal circulation and invades the fetus via the transplacental route. Bakardjiev et al. suggested a mechanism of transplacental infection through cell-to-cell spread from the decidua or from infected maternal immune cells, which subsequently infect the EVT (Bakardjiev et al., 2004; Bakardjiev et al., 2006). Once Lm enters the host cell, it can escape from the phagosomal vacuole via pores formed by the cytolysin listeriolysin O (LLO), replicate within the cytosol, and spread to neighboring cells by actin polymerization-mediated motility (Bakardjiev et al., 2005; Le Monnier et al., 2007). Invasive EVTs become accessible to maternal immune cells and thereby to Lm by the end of the first trimester (Knofler et al., 2019; Robbins et al., 2010). Lm is thought to traffic to the placenta within maternal immune cells without leaving this cellular “shuttle,” as has been observed with other pathogens like *Plasmodium falciparum*, which travels within erythrocytes to the placenta (Bakardjiev et al., 2006; Lecuit et al., 2004). Additionally, EVTs actively recruit maternal cells (Chaturvedi et al., 2015; Huang et al., 2008). Earlier experiments demonstrated that SYN acts as a barrier to Lm invasion and that Lm entry occurs in EVTs via cell-to-cell spread (Robbins et al., 2010). However, later work showed that EVTs restrict Lm infection (Zeldovich et al., 2011). In contrast, another transplacental invasion route was described by Lecuit et al. in placental explants. The authors found that placental infection can occur via direct invasion of Lm through the SYN layer in a receptor-mediated manner (Lecuit et al., 2004). The SYN express two key receptors, E-Cadherin (E-Cad) and c-Met (hepatocyte growth factor receptor), to which Lm can bind using two of its surface proteins Internalins, InlA and InlB, respectively (Lecuit, 2020). This was further substantiated by *in vitro* studies demonstrating that Lm infects trophoblast cell lines and human placental explants through InlA/E-Cad interactions. Immunohistochemical analysis of placental sections also revealed colocalization of Lm with SYN cells in regions with high E-Cad expression (Disson et al., 2008; Lecuit et al., 2004). Of note, these internalin-receptor interactions are species-specific, allowing Lm invasion in humans, guinea pigs, and gerbils, but not in mice due to a single amino acid difference in the mouse E-Cad at position 16 compared to human E-Cad (Lecuit et al., 1999; Disson et al., 2008; Dussurget et al., 2004). Consequently, this prevents InlA binding and thus the Lm placental infection risk in mice is low (Lecuit et al., 1999). To address this limitation, a humanized knock-in mouse strain, KIE16P, was generated, in which the glutamic acid at the 16th position of the mouse Ecad is replaced with a proline, as in human, allowing Lm infection in mice due to the functional InlA-Ecad interaction(Charlier et al., 2020; Disson et al., 2008). Moreover, internalins have also been used by Lm to cross other physiological barriers, such as the intestinal epithelial barrier (Lecuit et al., 2001), and the blood-brain barrier (Maudet et al., 2022). In the case of blood-brain barrier, Lm infected monocytes were identified to facilitate Lm crossing of this barrier aiding in subsequent neuroinvasion (Drevets et al., 2004; Join-Lambert et al., 2005; Maudet et al., 2022). Thus, while multiple mechanisms of cell-to-cell spread and direct invasion have been reported for Lm infection in the placenta, it remains unclear how Listeria reaches and crosses the maternal-fetal barrier during the early stages of infection.

In this study, we aimed to elucidate the mechanisms of early phases of placental and fetal infection by Lm, applying various *in vivo* and *in vitro* approaches, including the humanized E-Cadherin mouse KIE16P. We identified neutrophils as an intravascular shuttle and survival niche for Lm, facilitating Lm transfer and invasion of trophoblast cells in the placenta. *In vivo* experiments in KIE16P mice showed reduced placental and fetal Lm infection following neutrophil depletion or inhibition of neutrophil adhesion, confirming our *in vitro* findings, and highlighting a crucial role for neutrophils in Lm-mediated placental and fetal infection.

## Results

### Intravital observation of the mouse placenta shows immediate Lm clearance from the maternal circulation upon infection

To investigate the initial stages of Lm infection at the placenta and visualize potential maternal-to-fetal transmission *in vivo,* we established an intravital microscopy (IVM) model of the murine placenta (Fig. 1A-B). For these *in vivo* experiments, we used pregnant KIE16P mice (E13.5-15.5), a humanized mouse model for Lm infection (Disson et al., 2008). Using intravital multiphoton laser scanning microscopy, we observed Lm (injected via a carotid artery catheter) in the placental microvasculature. Interestingly, Lm was rapidly cleared from the maternal circulation, including the placental vasculature (Fig. 1C and Supplemental movie 1), aligning with previous findings on the rapid intravascular clearance of Lm in adult non-pregnant mice (Broadley et al., 2016; Verschoor et al., 2011). Furthermore, we found no sustained adhesion of Lm within placental microvessels, nor did we detect any transplacental transfer of Lm to the fetus (Supplemental movie 1). These findings suggest that Lm infection within the placenta is a rather rare event, challenging to be captured through intravital microscopy. Given the rapid clearance of Lm from the circulation, we hypothesized that the maternal circulation, particularly circulating blood cells, might act as intravascular carriers, providing a protective survival niche and a potential delivery system for shuttling Lm to the placenta.

**Figure 1.**
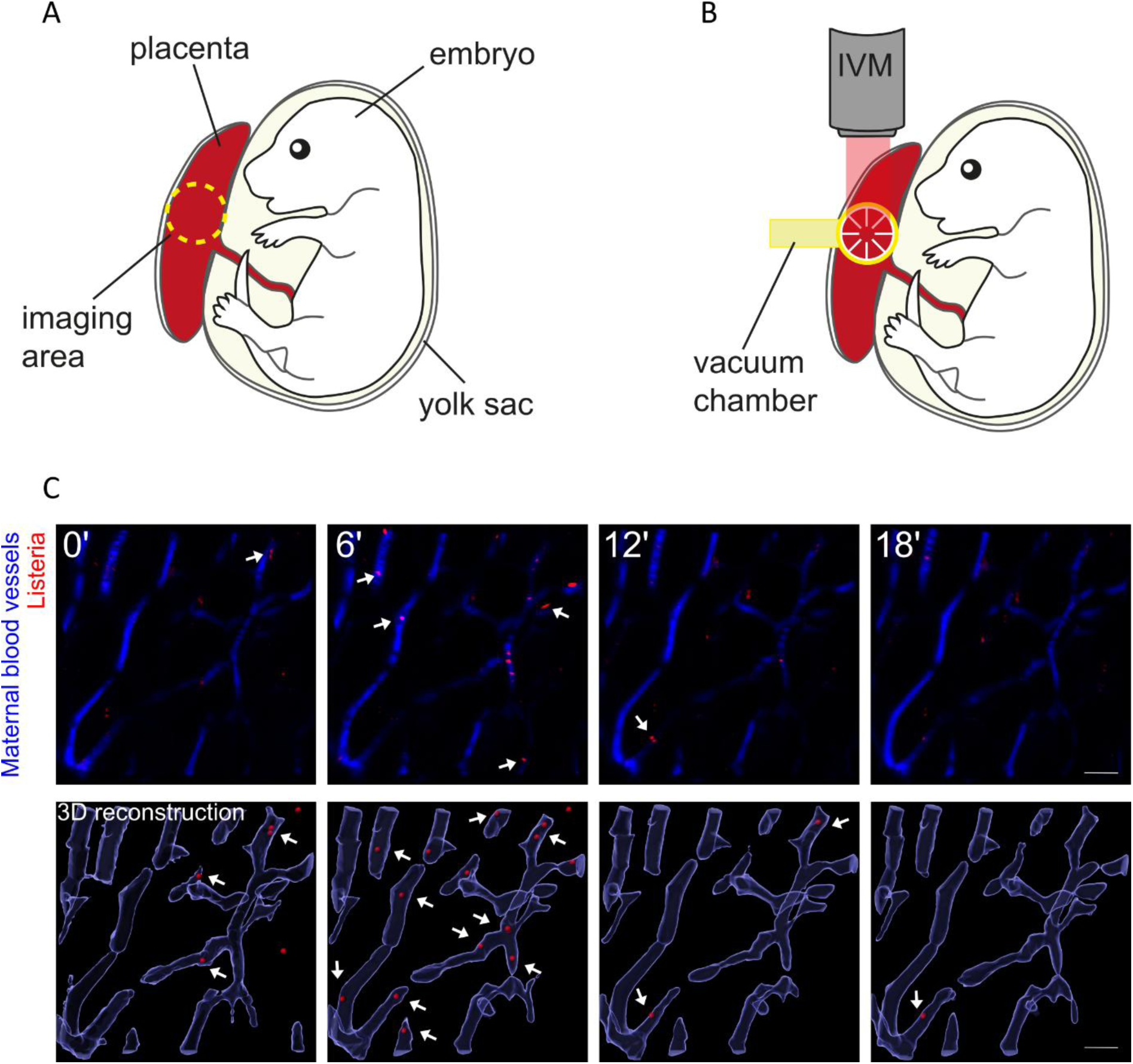
Intravital microscopy of Lm in the murine placental microcirculation. (A) Imaging area in the placenta (indicated by the yellow dotted circle) connected to the embryonic yolk sac. (B) Placement of the vacuum chamber holder for intravital imaging with positioning of the multiphoton objective for intravital imaging acquisition. IVM indicates intravital microscopy in the animation. (C) Transient appearance of Lm in maternal microvessels of the placenta visualized by multiphoton intravital microscopy and 3D reconstruction. Maternal blood labeled with blue fluorescent dextran dye, shows rapid clearance of Lm (red, upper panel). Time is displayed in minutes. White arrows indicate Lm (red). Scale bar = 20 µm.

### Human neutrophils serve as a survival niche and viability factor for Lm in the intravascular compartment

To test the hypothesis that Lm uses blood cells as shuttle and survival niche to reach the placenta, we collected whole blood from healthy human female volunteers, infected the blood with Lm for 1 h (MOI1) to study early infection windows, and subsequently treated the samples with gentamicin to eliminate all extracellular bacteria. Blood samples were stained with antibodies against CD14 (monocytes), CD56 (natural killer cells), and CD66b (neutrophils). CD41 binding to platelets was used as negative control, as Lm only binds to platelet surfaces (Broadley et al., 2016). Flow cytometry analysis of intracellular Lm in peripheral blood cells revealed that nearly 60% of neutrophils and monocytes carried intracellular bacteria, identifying them as a potential shuttle of Lm (Fig. 2A).

**Figure 2.**
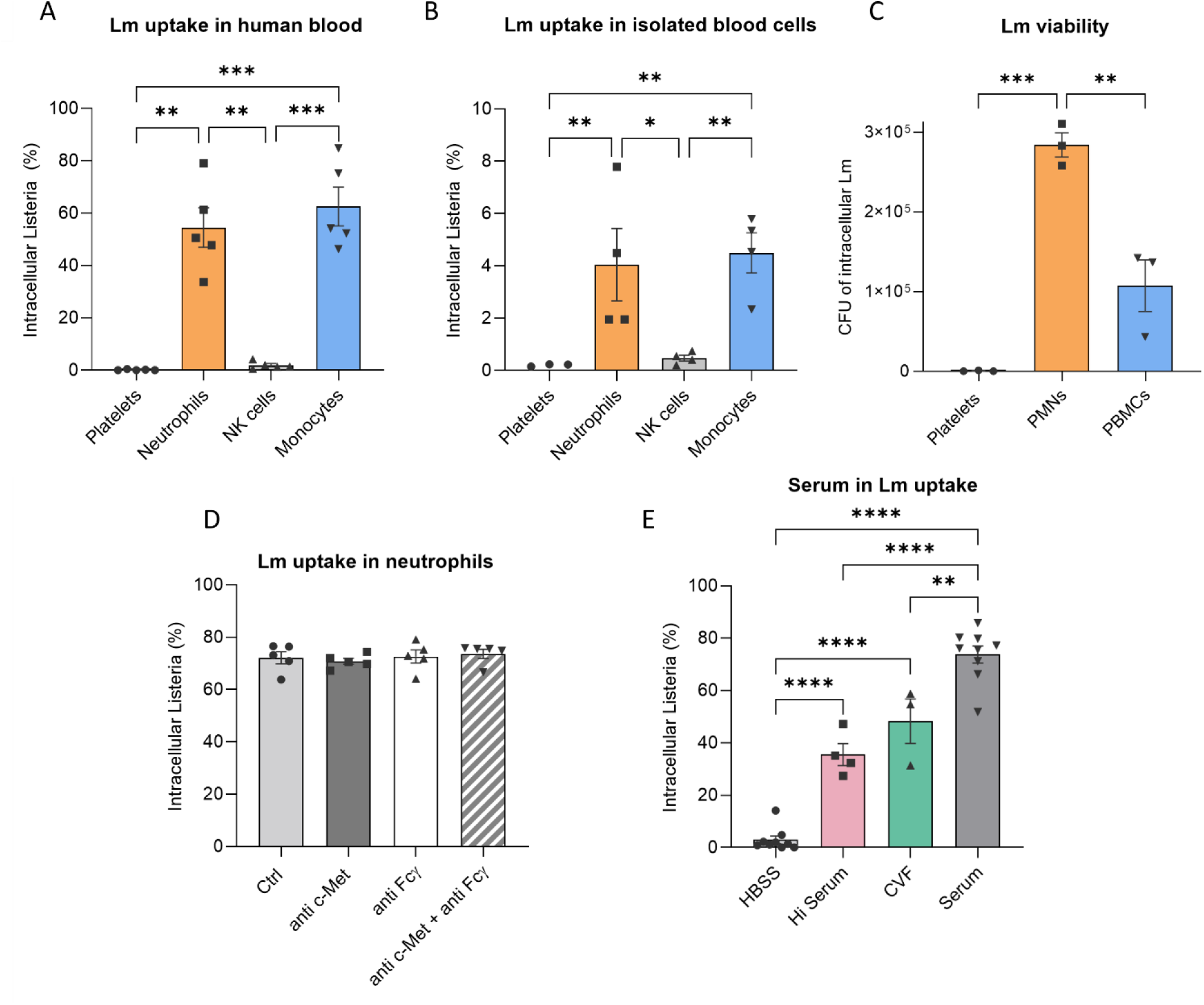
Human neutrophils serve as a survival niche and viability factor in the intravascular compartment. (A) Percentage of blood cells positive for CFSE-labeled intracellular Lm obtained from peripheral blood of non-pregnant human adults exposed to CFSE-labeled Lm (mean±SEM, n=5 independent experiments, 1-way ANOVA, Tukey’s multiple comparison). (B) Percentage of blood cells positive for CFSE-labeled intracellular Lm from isolated non-pregnant human peripheral blood cells exposed to CFSE-labeled Lm (mean±SEM, n=4 independent experiments, 1-way ANOVA, Tukey’s multiple comparison). (C) CFUs of intracellular Lm of Lm-infected lysed human neutrophils, PBMCs and platelets (mean±SEM, n=3 independent experiments, 1-way ANOVA, Tukey’s multiple comparison). (D) Percentage of human non-pregnant neutrophils positive for intracellular CFSE-labeled Lm after blocking c-Met and/or FcγR (mean±SEM, n=5 independent experiments, 1-way ANOVA, Tukey’s multiple comparison). (E) Percentage of human non-pregnant neutrophils positive for intracellular CFSE-labeled Lm after serum treatment via heat inactivation (Hi) and Cobra venom factor (CVF) treatment. Neutrophils in Hank’s Balanced Salt Solution (HBSS) without Lm-infection served as control (mean±SEM, n=3-11 independent experiments, 1-way ANOVA, Tukey’s multiple comparison). *p <0.05, **p <0.01, ***p<0.001, ****p<0.0001.

Next, we isolated neutrophils (PMNs, polymorphonuclear neutrophils), PBMCs (peripheral blood mononuclear cells), and platelets to investigate whether Lm remained viable within each cell type after uptake. After isolation, each cell population was infected with Lm (MOI8) and analyzed via flow cytometry. Similar to the whole blood findings, we identified neutrophils and monocytes as the primary carriers with the highest Lm load. Notably, the intracellular Lm content was much lower in isolated neutrophils (5%) as compared to neutrophils from whole blood infections (50% in Fig. 2A/B, Supplemental Figure 1), suggesting that serum components may enhance Lm uptake by neutrophils.

To determine whether Lm remains viable within these carriers/shuttle cells and whether Lm retains the potential to infect the placenta, we plated defined dilutions of infected isolated cells on selective antibiotic-supplemented agar plates (7 µg/ml) and counted colony-forming units (CFUs) the next day. Although neutrophils and monocytes had comparable levels of intracellular Lm, neutrophils harbored significantly more viable Lm (about 3-fold more) compared to PBMCs (Fig. 2C). This indicates that monocytes may be more effective than neutrophils at killing intracellular Lm, further suggesting a potential role of neutrophils as shuttle and viability factor for Lm to survive in the maternal intravascular compartment.

Given the significant viability of Lm in human neutrophils and that a potential serum factor(s) might facilitate Lm uptake, we further investigated this possibility in more detail. In addition, Lm uptake by monocyte-derived dendritic cells (MoDCs) occurs via opsonization with immunoglobulins and FcγRIII receptor-mediated internalization (Bortolussi et al., 1986; Kolb-Maurer et al., 2001). Another potential receptor, c-Met, which is also expressed on neutrophils (Finisguerra et al., 2015), was described to bind to InlB on Lm (Glodde et al., 2017). Furthermore, the complement receptor CR3 mediates Lm uptake by murine inflammatory macrophages (Drevets and Campbell, 1991). To examine whether these factors facilitate Lm uptake in human neutrophils, we used different antibody combinations and the cobra venom factor (CVF), which consumes C3, to block possible Lm uptake pathways into neutrophils. We found no effect of c-Met or FcγR on Lm uptake by human neutrophils (Fig. 2D). However, CVF-treated neutrophils showed reduced uptake of Lm by human neutrophils indicating that complement factor C3 contributes to Lm uptake by neutrophils (Fig. 2E). Interestingly, the stronger inhibition of Lm uptake observed in serum-treated samples compared to CVF treatment suggests that additional serum factors may facilitate Lm uptake by neutrophils (Fig. 2E).

### Lm evasion into neutrophils compared to monocytes

To further investigate the viability mechanism of Lm in neutrophils, we analyzed the intracellular location of Lm in neutrophils and monocytes post-internalization, hypothesizing that Lm may escape from vacuoles into the cytoplasm more effectively in neutrophils. For this, isolated neutrophils and monocytes were infected with pre-opsonized Lm for 1 hour at 37°C and then fixed overnight at 4°C. Transmission electron microscopy (TEM) analysis revealed significantly more Lm in the cytoplasm of neutrophils compared to monocytes (Fig. 3A-C), suggesting that Lm escape into the cytoplasm is more frequent in neutrophils than in monocytes, possibly aiding in Lm survival and subsequent infection.

**Figure 3.**
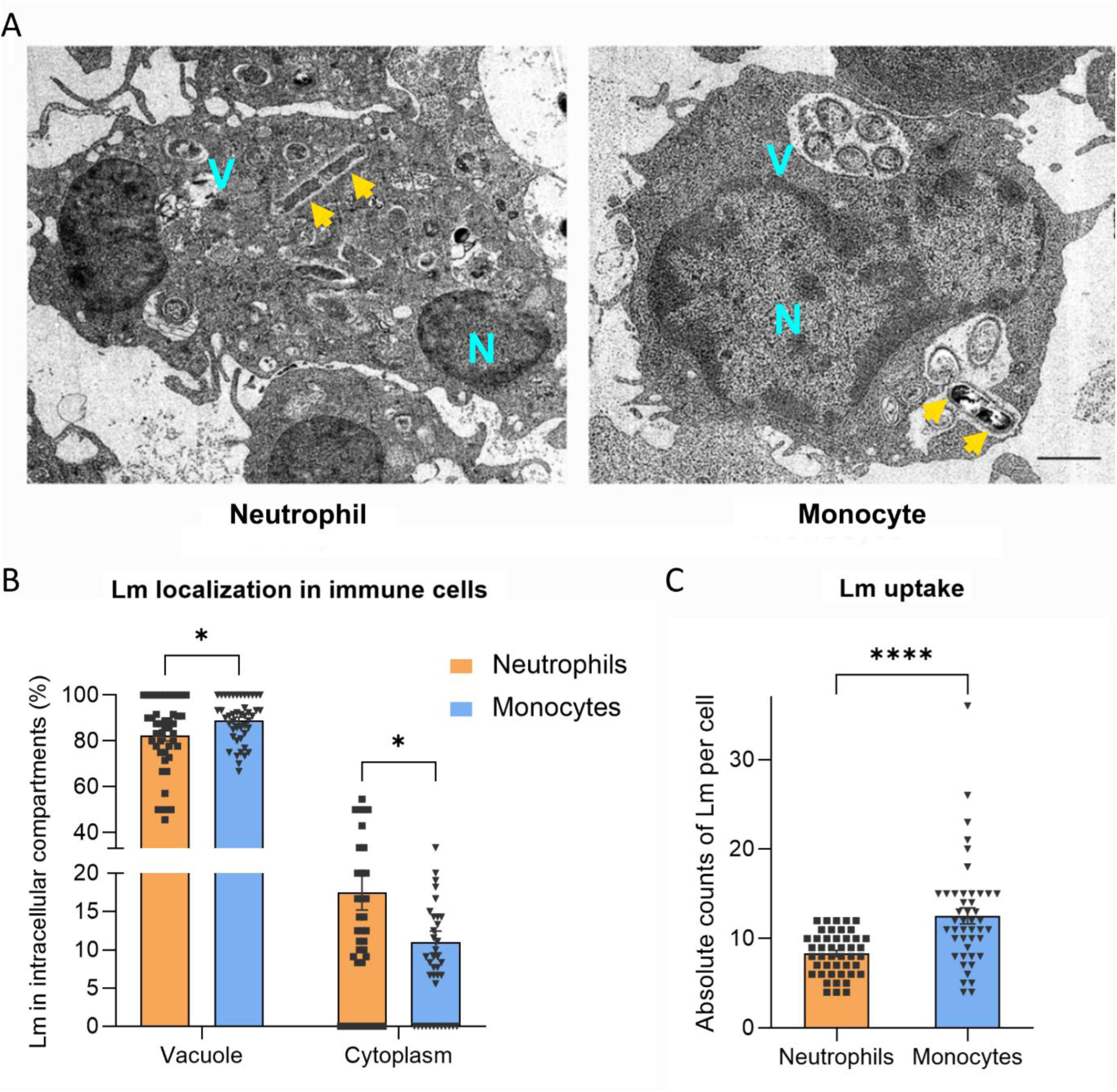
Cytoplasmic escape of Lm in neutrophils compared to monocytes. (A) Representative transmission electron microscopy image of an Lm-infected neutrophil and Lm-infected monocyte illustrating cytoplasmic versus vacuolar location of Lm. Listeria are represented with yellow arrows and V indicates vacuoles and N for nucleus. Scale bar = 1 µm. (B) Percentage of Lm in the vacuole and cytoplasm of both neutrophils and monocytes. (C). Absolute counts of Lm uptake in neutrophils and monocytes. Data shown as mean±SEM, n=45 cells/group, unpaired student‘s t-test; *p <0.05, ****p<0.0001.

### HTR8 trophoblast cells express adhesion relevant molecules

After collecting evidence that human neutrophils might serve as a survival niche for Lm, we next addressed potential mechanisms of how neutrophils successfully shuttle Lm to the placenta and mediate infection of the placenta and fetus. For that purpose, we chose two different human trophoblast cell lines, HTR8 and JEG3 cells. HTR8 cells, derived from extravillous trophoblast cells (EVT), were used to mimic a potential infection route through entering fetal tissue via EVTs. JEG3 cells are cytotrophoblast cells that can form syncytiotrophoblast cells (SYN) and were included to test whether their high levels of E-Cadherin might facilitate direct infection by Lm. Both trophoblast cell lines were investigated by flow cytometry and tested for their expression of adhesion relevant molecules as well as expression of E-Cad (InlA receptor) and c-Met (InlB receptor). On HTR8 cells, we found pronounced expression of ICAM-1, a critical adhesion molecule and ligand for neutrophil expressed β2 integrins LFA-1 and Mac-1. In addition, we found low expression of VCAM-1, PECAM-1, and E-selectin, but no detectable expression of P-Selectin, E-Cad, and c-Met relative to isotype control (Supplemental Figure 2A). As we could not detect E-Cad or c-Met expression via flow cytometry, we applied immunofluorescence staining and confocal microscopy and found c-Met expression, and faint E-Cadherin expression (Supplemental Figure 2B). For JEG3 cells, we found low expression of all adhesion relevant molecules tested including E-Cad and c-Met expression (Supplemental Figure 2C). Expression of E-cad and c-Met was also confirmed by confocal microscopy (Supplemental Figure 2D).

### Neutrophils mediate the invasion of Lm into TLR1/2-stimulated HTR8 trophoblast cells

So far, we identified human neutrophils as a potential carrier and viability niche for Lm in the intravascular compartment and could detect expression of neutrophil adhesion relevant molecules on trophoblast cells. Thus, we wanted to investigate whether neutrophils facilitate Lm entry into trophoblast cells of the placenta, initiating placental infection and transfer to the fetus. Therefore, trophoblast cells were analyzed concerning their vulnerability to Lm invasion. First, unstimulated HTR8 cells were infected with free Lm, Lm associated with platelets or Lm-infected neutrophils for 1 h. Thereafter, cells were lysed and plated on agar dishes to check for Lm viability by counting CFUs the next day. Alternatively, HTR8 cells were stained with antibodies against CD54 (ICAM-1) and analyzed via FACS to check for the intracellular Lm burden. We found no differences regarding amounts of intracellular Lm in HTR8 trophoblasts for all conditions compared to uninfected trophoblasts as control (Fig. 4A). Similarly, viability studies showed very low levels of CFUs (Fig. 4B).

**Figure 4.**
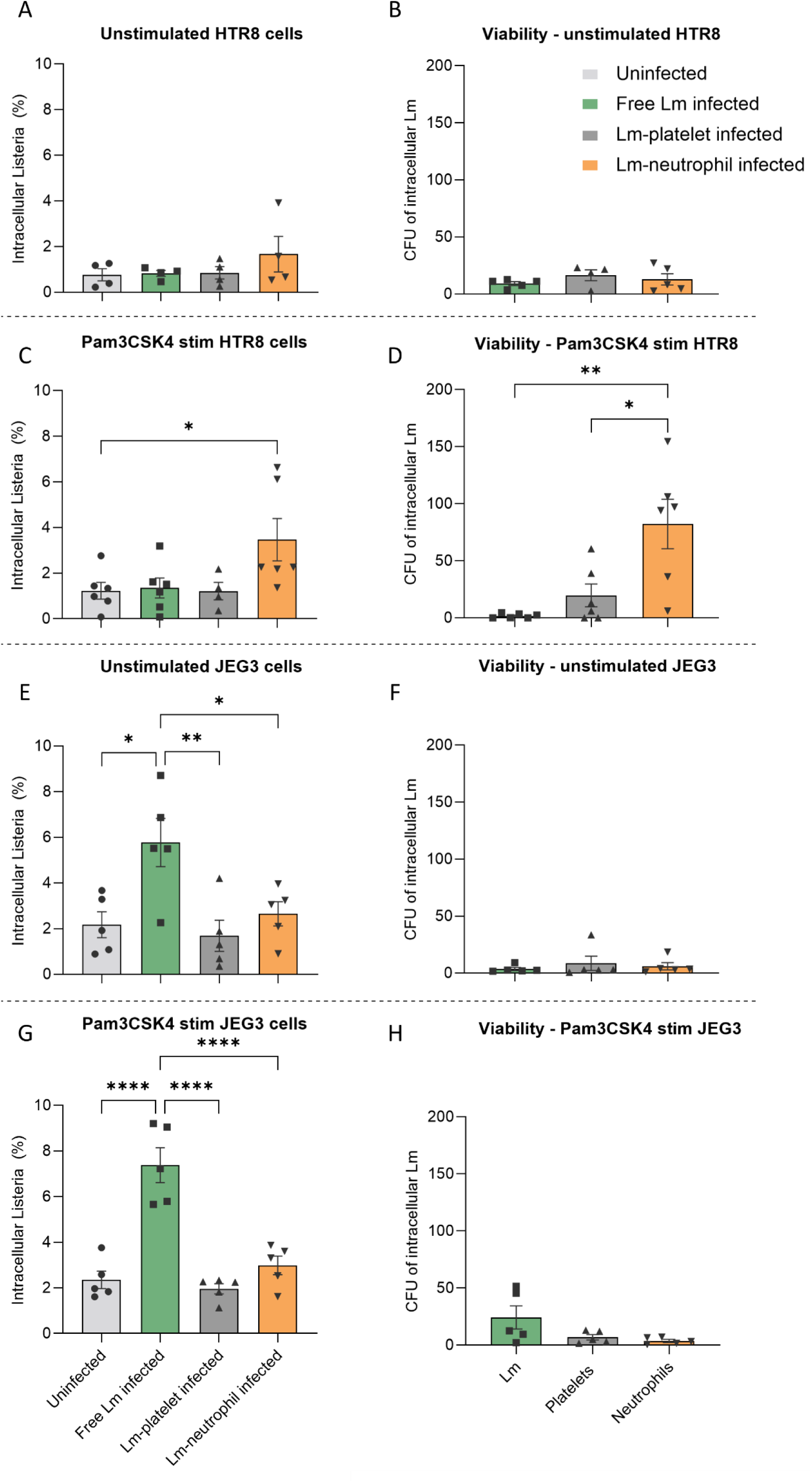
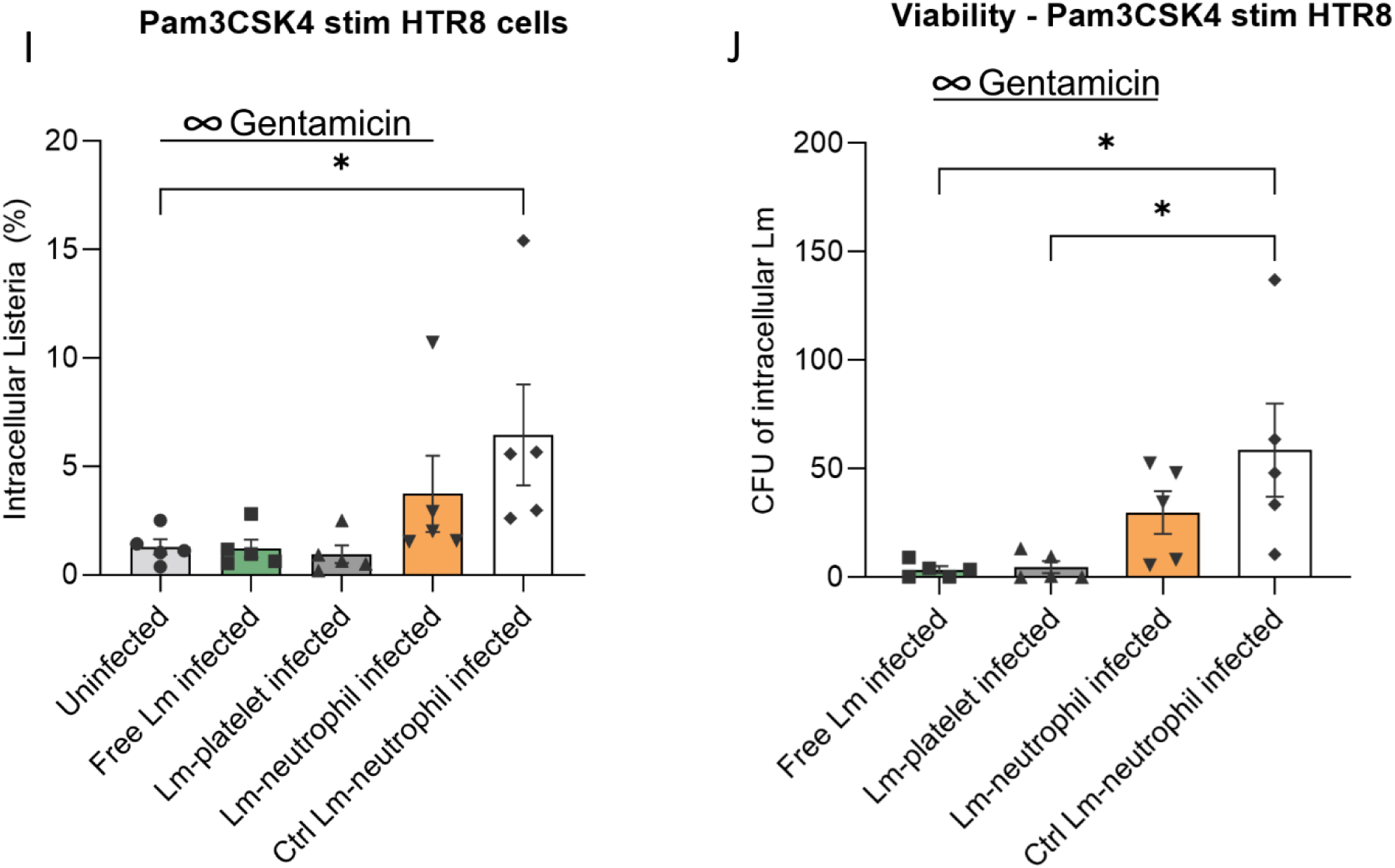
Invasion of HTR8 and JEG3 trophoblasts via free Lm, Lm-associated platelets and Lm-infected neutrophils. (A) FACS analysis and (B) viability check of Lm infection of unstimulated HTR8 cells (mean±SEM, n=4-5 independent experiments, 1-way ANOVA, Tukey’s multiple comparison). (C) FACS analysis and (D) viability check of Lm infection of Pam3CSK4-stimulated HTR8 cells (mean±SEM, n=4-6 independent experiments, 1-way ANOVA, Tukey’s multiple comparison). (E) FACS analysis and (F) viability check of Lm infection of unstimulated JEG3 cells (mean±SEM, n=5 independent experiments, 1-way ANOVA, Tukey’s multiple comparison) (G) FACS analysis and (H) viability check of Lm infection of Pam3CSK4-stimulated JEG3 cells (mean±SEM, n=5 independent experiments, 1-way ANOVA, Tukey’s multiple comparison) (I) FACS analysis and (J) viability check of Lm infection of Pam3CSK4-stimulated HTR8 cells following continuous gentamicin treatment for the first 4 groups and Lm-neutrophil infection without continuous gentamicin treatment for the last group (mean±SEM, n=5 independent experiments, 1-way ANOVA, Tukey’s multiple comparison). *p <0.05, **p <0.01, ***p<0.0001.

Chung and colleagues previously reported that TLR1/2 stimulation in the placenta increases the susceptibility to Lm infection (Chung et al., 2014). This made us investigate a potential role of TLR1/2 on Lm invasion using TLR2 agonist Pam3CSK4. When infecting Pam3CSK4-stimulated HTR8 trophoblasts, the intracellular amounts of Lm significantly increased up to four-fold after Lm-infected neutrophils were applied onto HTR8 trophoblasts compared to uninfected, free Lm-infected and Lm-platelet infected HTR8 cells (Fig. 4C). Interestingly, viability studies showed an even higher increase, reaching about 80-fold more intracellular viable bacteria in HTR8 cells infected with Lm-bearing neutrophils as compared to infections with free Lm and Lm-associated platelets (Fig. 4D). Taken together, these findings show that TLR2 stimulation of HTR8 cells significantly enhances transfer of Lm from infected neutrophils into HTR8 cells and its viability within HTR8 cells. This suggested that infection of the placenta by Lm may be facilitated in areas of the placenta with local inflammation and the presence of Lm-infected neutrophils.

Next, we investigated the susceptibility of JEG3 cells for infection by Lm, Lm-associated platelets or Lm-infected neutrophils. Interestingly, free Lm showed the best invasion of JEG3 compared to the other groups (Fig. 4E). However, intracellular Lm survival was rather low and similar to the other groups, suggesting that JEG3 cells effectively restrict intracellular Lm survival (Fig. 4F). When we repeated the same experiments with Pam3CSK4-stimulated JEG3 cells, we could neither observe a difference regarding the intracellular Lm burden (Fig. 4G) nor a change in the low numbers of CFUs counted on the day after the experiment (Fig. 4H). Aligning with previous reports (Zeldovich et al., 2011), this suggested a placental cell type specificity in their susceptibility to Lm infection and survival.

To further elucidate whether Lm enters trophoblast cells from maternal neutrophils via cell-to-cell spread or are released by neutrophils before they directly invade the trophoblast cells, gentamicin treatment was already started when free Lm, Lm associated with platelets or Lm-infected neutrophils were incubated with Pam3CSK4-stimulated HTR8 trophoblasts. This procedure eliminates the possibility that Lm released into the extracellular space by neutrophils bound to HTR8, can in turn invade HTR8 cells from the extracellular milieu (direct invasion resulting from neutrophil-mediated infection). To exclude any effects of gentamicin on neutrophil viability, we performed viability studies with neutrophils and found no significant cell death with gentamicin treatment for up to 4h (Supplemental Figure 3). Flow cytometric analysis revealed a partial reduction of Lm positive-HTR8 cells after incubation with Lm-infected neutrophils in the presence of extracellular gentamicin suggesting that Lm enters Pam3CSK4-stimulated HTR8 trophoblasts in part via E-cadherin dependent mechanisms (Fig. 4I). Viability studies confirmed the viability of transferred Lm, either through cell-to-cell spread or directly (Fig. 4J). Taken together, our results indicate that Lm can invade TLR2-stimulated HTR8 cells via cell-to-cell transfer from infected neutrophils or directly after their release from infected neutrophils.

### Mouse neutrophils internalize Listeria and allow their survival during pregnancy *in vivo*

Following the trophoblast experiments, we next asked if Lm-mediated placental infection occurs *in vivo* via neutrophils as well. To test whether Lm are taken up by immune cells in mouse peripheral blood, we injected CFSE-labeled Lm (EGDe-GFP) into non-pregnant and pregnant (E13.5-14.5) C57BL/6 mice via intravenous (i.v.) injection. The temporal kinetics of Lm viability in whole blood were tested by CFU plating across various time points of infection based on previous reports (Broadley et al., 2016; Koopmans et al., 2018; Verschoor et al., 2011; Wang et al., 2011) (Supplemental Figure 4A). Accordingly, 15 min after Lm infection was selected for assessing Lm-immune cell dynamics for further experiments.

Subsequently, whole blood was harvested from Lm or mock-infected pregnant/non-pregnant mice, followed by gentamicin treatment and staining for antibodies against CD45 (leukocytes), CD3 (T-cells), CD19 (B-cells), Ly6G (neutrophils) and Ly6C (monocytes) (Supplemental Figure 4B). FACS analysis showed no significant changes in immune cell distribution of the various mouse groups (Supplemental Figure 4C, D). Interestingly, Lm uptake was predominantly observed in Ly6G+ neutrophils (Fig. 5A), while much reduced uptake was seen in other immune cell populations, including CD11b+ Ly6C^high^ monocytes (Supplemental Figure 5A, B). Notably, in contrast to monocytes, neutrophils from pregnant mice had an elevated Lm uptake as compared to non-pregnant mice (see Fig. 5A, B). CFU counts of intracellular Lm from whole blood of pregnant mice also showed higher Lm viability (Fig. 5C). Since the majority of Lm uptake via FACS was seen in Ly6G+ neutrophils, it can be correlated at the group level to the higher viability observed via CFU counts. This was in agreement with the Lm dynamics of ex vivo human blood experiments as well.

**Figure 5.**
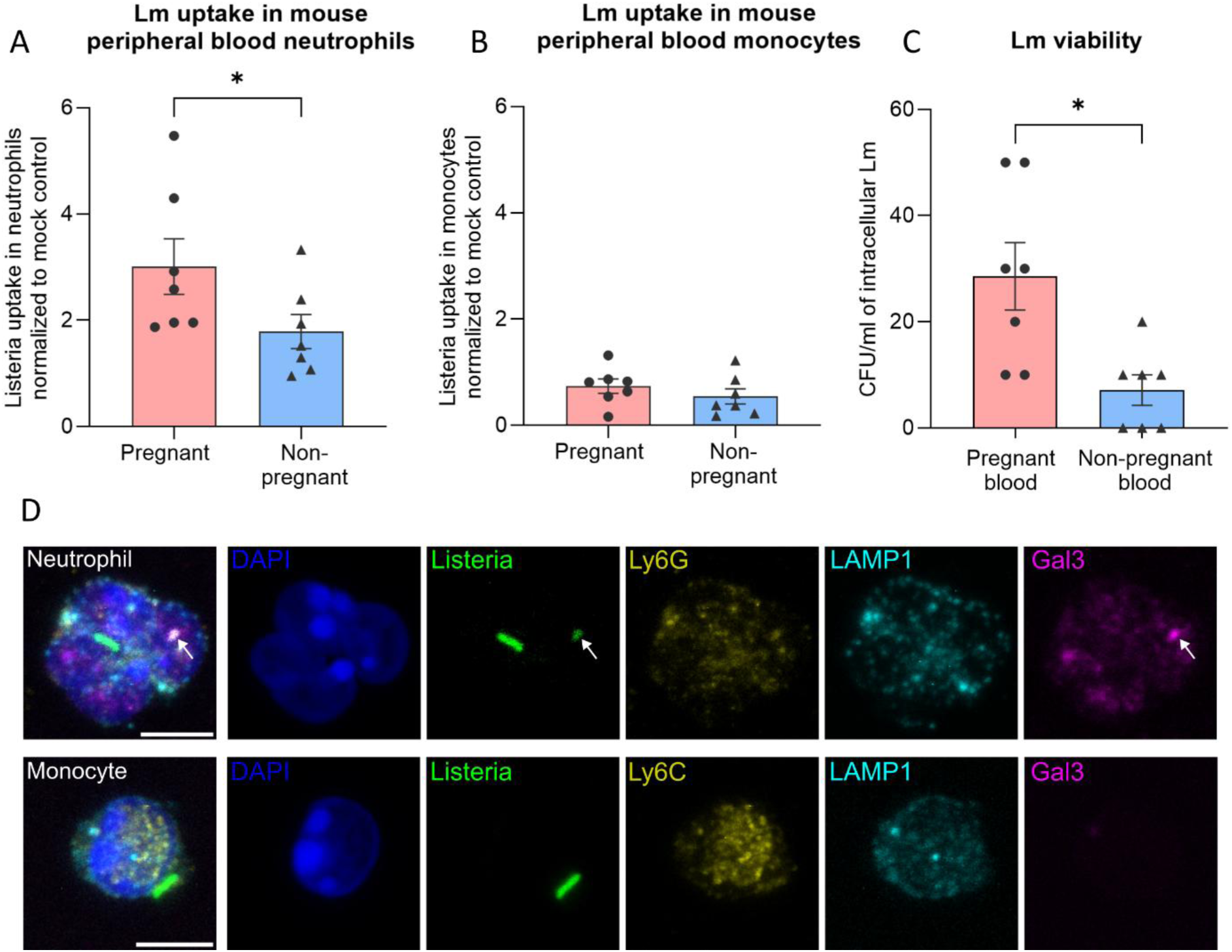
Pregnant mouse neutrophils harbor viable Listeria in the intravascular compartment. (A-B) FACS analysis of *in vivo* Lm infection in peripheral blood of pregnant and non-pregnant mice neutrophils (A) and monocytes (B). Data are shown for CD45+ CD3-CD19-Ly6G+ neutrophils (A) and CD45+ CD3-CD19-Ly6G-CD11b+ Ly6C+ monocytes (B). CFSE-positive Lm uptake in immune cells of infected mice is normalized to corresponding background signal in mock-infected controls (mean ± SEM, n=7 mice per condition, unpaired student’s t-test). (C) Intracellular viable bacteria in Lm-infected pregnant and non-pregnant mouse blood were analyzed by CFU analysis of gentamicin-treated and lysed blood samples, shown as CFU per ml of blood (mean ± SEM, n=7 mice per condition, unpaired student’s t-test with Welch’s correction). (D) Confocal microscopy imaging of Lm-infected pregnant mouse leukocytes. Top panel shows the maximum intensity projection of an infected pregnant mouse neutrophil identified by the DNA staining (DAPI shown in blue) and neutrophil-specific membrane marker (Ly6G shown in yellow), with Listeria inside (green). Internalized Listeria colocalizes with lysosomal escape marker protein, Galectin-3 (magenta) and does not associate with lysosomal compartments (labelled with LAMP-1, lysosomal membrane protein in cyan). White arrows correspond to Lm associated with Galectin-3 punctae recruitment indicative of vacuolar escape and intracellular survival. Bottom panel shows an image of a monocyte identified by DAPI staining as well as monocyte-specific membrane marker (Ly6C in yellow). Internalized Lm is not associated with Gal3 signal, suggesting no cytosolic escape and potential survival. Scale bars = 5 µm. Images representative of 3-4 pregnant mice. *p <0.05.

To further corroborate that Lm utilizes maternal neutrophils as viability shuttles *in vivo*, we performed confocal microscopy imaging of leukocytes from Lm-infected pregnant mice. CFSE-positive Lm signal was detected within neutrophils and less frequently in monocytes, identified by their characteristic nuclear DNA morphology and membrane markers, Ly6G and Ly6C, respectively. To ascertain if Lm can survive in these phagocytic cells via vacuolar rupture, the localization of Lm within vacuoles (marked by LAMP-1) and ruptured vacuoles were examined. For this, we stained intracellular Galectin-3 (Gal-3) to mark early phagocytic vacuolar rupture and thus cytosolic escape of Lm (Aits et al., 2015; Jia et al., 2020; Paz et al., 2010; Weng et al., 2018). Strikingly, we observed a colocalization of internalized Listeria with Gal-3 punctae in neutrophils, consistent with phagosomal damage, but not in monocytes that had taken up Lm (Fig. 5C, Supplemental Figure 5C, E). Accordingly, Listeria that were confined to lysosomes, did not recruit Gal-3 (Supplemental Figure 5C-E). Taken together, these results suggest that Lm infects and remains viable in neutrophils in the maternal circulation, via escape into the cytoplasm, at early time points, to potentially mediate placental infection *in vivo*.

### Neutrophils colocalize with Lm in the placenta

To investigate whether neutrophils are indeed associated with Lm in the placenta upon early infection, we used a tissue clearing approach of Lm-infected placentas. Following intraarterial injection of GFP-labeled Lm (EGDe-GFP) into pregnant KIE16P mice (E12.5-15.5), mice were sacrificed 1h after injection of Lm. After killing the mice, placentas were collected and prepared for whole organ clearing. 3D reconstruction of the placentas revealed the spatial distribution of Lm and neutrophils with detection of colocalized Lm with neutrophils (Fig. 6A, Supplemental Movie 2). To quantify the colocalization rate, we used a deep learning model (https://github.com/erturklab/SCP-Nano) to segment the bright dots representing Lm in the images. We then performed statistical analysis on the intensity values to determine an appropriate threshold for segmenting the bright regions representing neutrophils (Fig. 6B). By calculating the ratio of segmented Lm within neutrophil regions to the total segmented Lm, we found the colocalization rate of Lm and neutrophils ranged from 2.94% to 17.69% across four placentas (Fig. 6C). These findings indicate colocalization of Lm and neutrophils in the placenta implying a role of maternal neutrophils carrying Lm in mediating infection across the maternal-fetal barrier.

**Figure 6.**
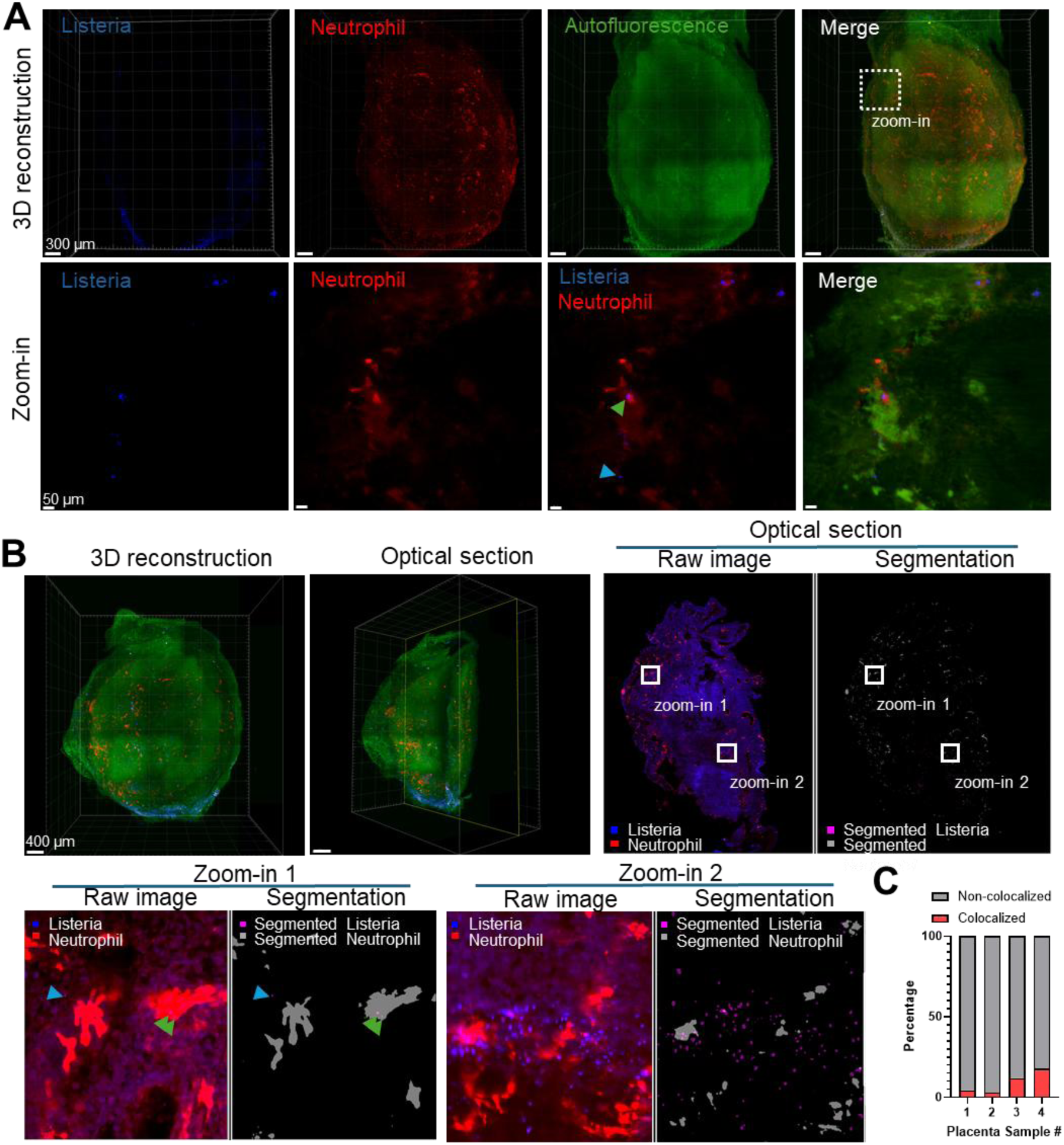
Visualization and colocalization analysis of Lm and neutrophils in intact cleared placentas. (A) 3D reconstruction of whole mouse placentas with Lm in blue, neutrophils in red, and tissue auto-fluorescence in green. Green arrowhead indicates areas where Lm and neutrophils colocalize, while blue arrowheads indicate Lm outside neutrophil regions. (B) Overview of the colocalization analysis pipeline. Green arrowheads mark colocalized Lm and neutrophils, and blue arrowheads mark Lm outside neutrophil regions. Scale bars are indicated on the images. (C). Quantification of Lm instances colocalized and non-colocalized with neutrophils across 4 placentas.

### Neutrophils are critical for placental und fetal infection

After having shown that maternal neutrophils provide a niche for Lm in the intravascular compartment and colocalize with Lm in the infected placenta, we aimed to confirm and strengthen our findings on the role of neutrophils during Lm infection of the placenta and fetus under *in vivo* conditions. To do this, we used pregnant KIE16P and wild type (WT) BL/6 mice (E13.5-E14.5) and depleted neutrophils by injection of anti-Ly6G antibody i.v. 24 h and then 4 h before Lm was injected i.v. (Supplemental Figure 6A). Eight hours after Lm injection (selected as an early time point for Lm-mediated tissue colonization and within the lifetime of circulating neutrophils (Koopmans et al., 2018)), mice were sacrificed, and placentas, fetuses and maternal liver (control organ) taken out, homogenized and cell suspensions plated on agar dishes to count CFUs and thus determine the intracellular bacterial burden/load in the respective organs. In a second approach, we injected antibodies against the β2 integrins CD11b (MAC-1) and CD11a (LFA-1) i.p. 2 h before i.v. injection of Lm to prevent neutrophil attachment to inflamed microvessels (Wen et al., 2022). Eight hours later, mice were sacrificed and organs processed as described above. When analyzing the amount of viable Lm in placentas (Fig. 7A) and fetuses (Fig. 7B, Supplemental Figure 6B), we saw a significant decrease of bacterial burden in the placentas of KIE16P mice after neutrophil depletion or after blocking neutrophil adhesion compared to untreated conditions, whereas it had no effect on bacterial load in C57BL/6 mice, which was already low without neutrophil depletion (Fig. 7A), in accordance with previous reports (Poulsen et al., 2011). For the percentage of infected fetuses, we could observe a similar pattern, showing a decrease of fetal infection in KIE16P mice from above 30 % to 10 % after blocking neutrophil adhesion or neutrophil depletion and surprisingly no infection of WT mice at all when neutrophils were depleted (Fig. 7B, Supplemental Figure 6C). Neutrophils are known to contribute to early clearance of Lm from tissues such as liver and spleen, at time points similar to those used in our experiments (Carr et al., 2011). Prior studies have also shown that the genetic background plays a role, with non-C57BL/6 mice exhibiting increased susceptibility to hepatic Lm infection, likely due to reduced neutrophil-mediated clearance (Pitts et al., 2018). While a slight reduction in liver CFU counts were observed in KIE16P mice depleted of neutrophils, this trend was not statistically significant and maternal liver CFU counts remained unchanged across groups (Fig. 7C), highlighting neutrophil-mediated specificity for placental and fetal infection.

**Figure 7.**
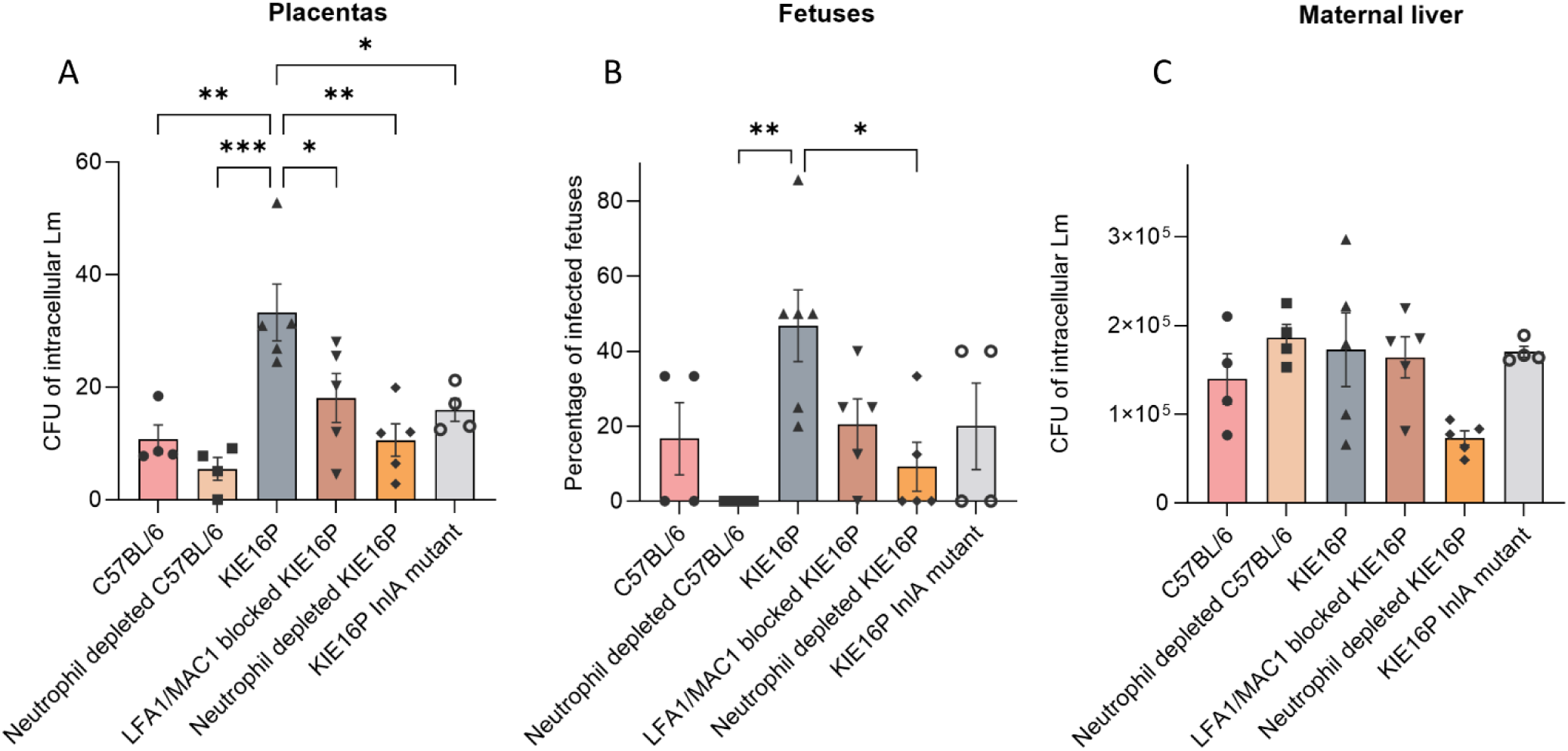
Lm infection of placentas, fetuses, and maternal livers. (A) Listeria load of placentas presented in amount of CFUs per mouse of various conditions. (B) Overall percentage of infected fetuses per condition. (C) Listeria load in maternal livers (control organ) presented as CFUs per mice. All data shown as mean±SEM, n=4-5 mice per condition, 1-way ANOVA, Tukey’s multiple comparison). *p <0.05, **p <0.01, ***p <0.001.

These findings clearly indicate that depletion of neutrophils or blocking adhesion of neutrophils impairs placental and fetal infection with Lm in KIE16P mice suggesting a role of E-Cadherin in this process. To confirm a role of human E-Cadherin expressed in KIE16P (but not in WT mice), we next infected pregnant KIE16P mice with the InlA deletion mutant of the EGDe strain. We observed a significant decrease of the bacterial burden with the InlA deletion Lm mutant in placentas and fetuses of KIE16P mice, which was similar to results seen after neutrophil depletion or blocking of neutrophil adhesion (Fig. 7A, B). Interestingly, maternal infection of the liver was rather independent of InlA and E-Cadherin (Fig. 7C). These findings clearly demonstrate a critical role of human E-Cadherin (in humanized KIE16P mice), following cell-cell transfer, in Lm infection of the placenta and fetus. Altogether, we show that Lm exploits maternal neutrophils in the circulation as transient viability reservoirs, facilitating infection across inflamed regions of the maternal-fetal barrier at early time points.

## Discussion

*Listeria monocytogenes* (Lm) infection can pose a serious risk to pregnant women and through transplacental transmission also to their growing fetus. In this study, we identify maternal neutrophils as a critical survival niche and vector for Lm within the maternal circulation, facilitating its transfer to placental tissue. Following maternal Lm infection, neutrophils harbor Lm intracellularly, protecting Lm from rapid clearance and enabling Lm’s delivery to inflamed trophoblast cells in the placenta, from where Lm spreads to the fetus. Although previous studies have highlighted placental susceptibility to Lm, the exact molecular mechanisms underlying placental invasion at early infection stages remained unclear.

Using intravital multiphoton microscopy, we extended findings of the rapid clearance of Lm from the maternal bloodstream, as observed in non-pregnant mice, to pregnant models (Broadley et al., 2016; Verschoor et al., 2011). Due to this rapid clearance mechanism, we could not directly observe Lm infection of the placenta by *in vivo* imaging. However, our tissue cleared-placenta imaging analyses, neutrophil depletion experiments as well as neutrophil adhesion blocking experiments *in vivo*, indicate successful placental infiltration of Lm occurring at least in part through a neutrophil-mediated transport. Recent studies indicate that bovine PMNs may allow intracellular survival of Lm for re-infection of other cell types, potentially to mediate bovine listeriosis (Bagatella et al., 2025). This mechanism has also been discussed for other intracellular pathogens that co-opt host immune cells to facilitate transport into the placenta (da Silva et al., 2024) supporting a model in which Lm evades immune defenses by hijacking leukocytes for transmission into placental tissue (Robbins et al., 2010). Of note, recent work from Maudet and colleagues showed that Lm can cross the blood-brain barrier by utilizing monocytes to avoid immune cell attack and promote infection (Drevets et al., 2004; Join-Lambert et al., 2005; Maudet et al., 2022). Accordingly, the identification of neutrophils as a shuttle, survival niche, and viability factor for Lm in the intravascular compartment was unexpected, as monocytes had been considered the primary host cell type for Lm, partly also due to clinical reports of mononuclear leukocytosis associated with Lm infection in mouse models (Cossart, 2007; Drevets et al., 2004; Maudet et al., 2022). Neutrophil depletion studies have further emphasized the indispensable role of monocytes in Lm liver infections, suggesting that monocytes, rather than neutrophils, are crucial for the spread of Lm in hepatic tissue (Witter et al., 2016). Similarly, studies in the gut have shown that Lm-infected monocytes exhibit the highest bacterial load, but only few bacteria manage to reach the cytosol, limiting Lm replication and survival (Jones and D’Orazio, 2017) which might indicate that other leukocyte subsets including neutrophils might also contribute to Lm infection. Accordingly, studies on how Lm disseminates from the gut to other tissues, such as lymph nodes, show that monocytes are not required for the colonization of lymph nodes and that multiple mechanisms may be used for Lm spread from the gut (Nowacki et al., 2025). Interestingly for placental Lm infection, monocyte-trophoblast interactions in the placenta are shown to trigger inflammasome activation in monocytes, resulting in the effective killing of intracellular Lm, highlighting the robust defense mechanisms that monocytes possess to limit Lm spread to the placenta (Megli et al., 2021). This supports the idea that other leukocytes such as neutrophils, might have a role in placental Lm infection. Our own experiments corroborate this and show that neutrophils are able to internalize substantial amounts of Lm and retain them in the cytoplasmic compartment while circulating in blood. The ex vivo human blood and *in vivo* mouse blood experiments of Lm infection show that these are early-stage Lm responses that allow them to potentially persist in the circulation avoiding antimicrobial attack.

In the blood circulation, Lm appears to survive within neutrophils following uptake, a process partially facilitated in the vasculature by complement factor C3. Lm has developed multiple adaptive strategies to support host survival and persistence, which are especially effective within neutrophils. One of the major virulence factors of Lm, listeriolysin O (LLO), enables vacuolar escape to the cytoplasm and becomes active at a pH of 5.5–6.0, the acidic range found in early phagosomes (Dussurget et al., 2004; Nguyen et al., 2019; Westman and Grinstein, 2020). Additionally, neutrophils exhibit a higher phagosomal pH compared to monocytes or macrophages (Foote et al., 2019; Nordenfelt and Tapper, 2011), potentially facilitating Lm’s cytoplasmic escape. Another reason for the intracellular persistence of Lm are bacterial enzymes like catalase and superoxide dismutase which help Lm to counteract oxidative bursts by neutralizing reactive oxygen species (ROS) (Pitts et al., 2018). This is further demonstrated by the reduced ROS production of neutrophils when exposed to Lm as compared to other bacteria, suggesting an inability of neutrophils to readily clear Lm (Richardson et al., 2024). All such protective adaptations enable Lm to evade neutrophils’ oxidative defenses, thereby enhancing their survival and replication within these cells.

However, differences in inflammasome activation between monocytes and neutrophils may also be of importance (Hybiske and Stephens, 2015). This is also highlighted in inflammasome-mediated Lm defense in monocyte-trophoblast interactions upon placental infection (Megli et al., 2021). Most bactericidal functions in neutrophils are localized within vacuoles, which is an intracellular compartmentalization also observed in other pathogens like *Burkholderia thailandensis*, suggesting that a similar mechanism may apply to Lm, at least temporarily (Kovacs et al., 2020). Interestingly, we recently showed that fetal/neonatal neutrophils exhibit reduced inflammasome activation as compared to adults (Wackerbarth et al., 2024). Further reports in pregnancy implicate inflammasome signaling in placental tissues and maternal compartments in healthy and complicated pregnancies, supporting the idea that pregnancy-specific inflammasome activation could modulate immune cell behaviour during infection (Gomez-Lopez et al., 2019; Moss et al., 2025). Generally, survival within phagocytes, particularly neutrophils, is uncommon; however, our analysis identified significantly higher numbers of viable Lm in the cytoplasm of neutrophils (consistent with galectin-3 recruitment indicative of early phagosomal damage), as compared to PBMCs. These early-stage Lm escape-responses into the cytosol are potentially sufficient for them to persist in maternal neutrophils till they reach the placental barrier, following Lm dissemination into the placenta and causing fetal infection.

During pregnancy, an increase in the number of neutrophils (though not monocytes) is observed in humans throughout the stages of pregnancy, which may give Lm a quantitative advantage by providing more potential host cells for hiding and subsequent infection of the placenta and fetus (Bert et al., 2021; Gomez-Lopez et al., 2014). Neutrophils during pregnancy have also been shown to have altered functional profiles with responses to inflammatory stimuli to contribute to maternal-fetal tolerance and this can potentially modulate their defense against early Lm infection kinetics aiding Lm survival (Bert et al., 2021; Calo et al., 2025; Gimeno-Molina et al., 2022). In addition, labor-associated neutrophils are reported to be in an activated state, with an enhanced capacity for migration (Gomez-Lopez et al., 2014). Elevated mRNA levels of CXCL8, for example, have been observed in the myometrium and decidua, correlating with higher neutrophil numbers in these regions, especially under inflammatory or infectious conditions (Gomez-Lopez et al., 2014). Further, the migration of leukocytes into the placenta during processes such as spiral artery remodeling may further increase the risk of Lm infection. This risk is amplified by pro-inflammatory conditions that prevail until the end of the third trimester, a period when highly active decidual neutrophils support a pregnancy-sustaining environment, which could be opportunistically exploited by Lm (Giaglis et al., 2016; Racicot et al., 2014). Accordingly, neutrophil infiltration has been observed in placentas with Listeria pathology which were previously attributed to infection responses, but our study, for the first time, hints at a role of neutrophils in transferring Lm to the placenta (Fell et al., 2025; Le Monnier et al., 2006; Richardson et al., 2024).

Concerning transfer of Lm from neutrophils to trophoblast cells, our findings demonstrate that Lm-infected neutrophils facilitate Lm invasion into the extravillous trophoblast cell line HTR8 only following TLR2 stimulation, while the cytotrophoblast cell line JEG3 is susceptible to direct Lm infection without requiring stimulation, suggesting a cell type and cell status-depending Lm specificity. Although JEG3 cells allow initial invasion by free Lm, our viability assays show that they effectively restrict Lm’s intracellular growth and spread. This suggests that placental invasion by Lm through infected cytotrophoblast cells might be a more uncommon event.

In contrast, we observed a significant increase in viable intracellular *Listeria monocytogenes* within HTR8 cells when Lm-infected neutrophils were co-cultured with them. This finding aligns with previous reports, such as those by Robbins and colleagues, who demonstrated that syncytiotrophoblasts (SYNs), which are in constant contact with maternal blood, exhibit strong resistance to bacterial spread (Johnson et al., 2021; Robbins et al., 2010; Zeldovich et al., 2011). Similarly, Megli et al. observed a high level of resistance to Lm infection in midgestation chorionic villi, with little to no bacterial growth—a pattern consistent with our *in vitro* findings using JEG3 cells (Megli et al., 2021).

The negligible role of SYN in placental Lm infection, as proposed here, is further supported by recent studies that highlight the binding of Lm internalin P (InlP) to afadin, a scaffolding protein associated with cell-cell junctions. This interaction disrupts cellular integrity, facilitating Lm transcytosis across the basement membrane and ultimately enabling fetal infection (Faralla et al., 2016). SYN have been shown to exhibit strong resistance to Lm infection due to their unique cellular structure, which lacks cell-cell junctions—and thus afadin—preventing InlP-mediated attachment and limiting fetal spread (Ander et al., 2019). In contrast, extravillous trophoblasts (EVTs) do possess cell junctions with afadin, allowing InlP-afadin interactions to support Lm invasion and spread in these cells (Faralla et al., 2018). This mechanism likely contributes to the resistance observed in JEG3 cells in our experiments, as InlP-afadin interactions are absent in SYN-like cells, limiting Lm’s capacity for intracellular growth and spread in this trophoblast subtype (Faralla et al., 2018). These studies further highlight the importance of placental tropism in propagating Lm infection, suggesting that Lm survival in neutrophils is associated with specific placental cell niches.

To investigate whether Lm enters trophoblasts cells via cell-to-cell spread or direct invasion, we conducted experiments with continuous gentamicin treatment. Our findings demonstrated that Lm can enter TLR2 agonist Pam3CSK4-stimulated HTR8 trophoblasts through both mechanisms: cell-to-cell spread and direct transfer into the cells following release from or exit out of neutrophils. This is also in line with the observations of free-floating and cell-associated Lm in invading other tissues from the gut (Nowacki et al., 2025). Pathogen release from host cells has been described to be occurring via both lytic or non-lytic processes (Hybiske and Stephens, 2015). One such exit mechanism, pyroptosis, primarily functions as a host defense strategy against intracellular pathogens like Lm (Flieger et al., 2018). Lm has been shown to utilize pyroptosis as an exit strategy in endothelial cells and macrophages, suggesting a similar process might occur in neutrophils (Hybiske and Stephens, 2015). However, recent work from our group and others indicated that neutrophils exhibit considerable resistance to pyroptosis (Chauhan et al., 2022; Pruenster et al., 2023). This resistance may allow neutrophils to serve as stable reservoirs for Lm, facilitating its transfer to trophoblast cells in placental tissues.

To further substantiate the critical role of neutrophils in the entry of Lm into trophoblast cells leading to subsequent placental infection, we translated our in vitro findings to experiments under *in vivo* conditions. Here, neutrophils in peripheral blood showed an increased Lm uptake, specifically in pregnant mice, with enhanced intracellular survival, as observed via the lysosomal escape microscopy imaging as well. We propose that placental Listeria infection requires maternal neutrophils as an intravascular shuttle to provide spatial access for Lm to the placental cells. The reported low shear stress profiles of maternal blood within placental microvessels (Jensen and Chernyavsky, 2019; Lecarpentier et al., 2016), potentially favors this by aiding the neutrophils carrying Lm to be possibly stalled in the placenta for infection. Accordingly, blocking neutrophil adhesion molecules LFA1 and Mac-1, as well as depleting neutrophils from the maternal circulation, resulted in impaired placental and fetal infection, particularly in KIE16P mice. This finding also underscores the significant role of human E-Cadherin in this process. The importance of E-Cadherin was further supported by observations of reduced placental and fetal infection of KIE16P mice infected with an Lm mutant defective in the InlA gene. Additionally, we performed whole organ clearing of the placenta to detect colocalization of neutrophils with Lm. We identified colocalization of sites, although they were rare in number, in par with the early time points of infection used here. Adding onto this, Lm can induce brain infections with rare invasion events at early stages of infection and can seed the placenta with a single bacterium to cause placental infections (Bakardjiev et al., 2006; Chevee et al., 2024). This result was thus not unexpected, and along with our intravital microscopy approach and *in vitro* trophoblast infection experiments indicated that placental infection is a rare event, most likely facilitated by locally inflamed trophoblast cells.

Our studies highlight that maternal neutrophils in the circulation during pregnancy are repurposed by Listeria to propagate placentofetal invasion at early stages of infection. Following this, it is plausible that the later stages of infection are associated with downstream immune cell recruitment, as reported for neuroinvasion with inflammatory monocytes (Maudet et al., 2022) or for placental invasion with CD8+ T-cells causing fetal loss (Chaturvedi et al., 2015). The placenta itself provides multiple defenses against Lm, which may be partially hindered by the association of Lm within neutrophils initially (Megli et al., 2021). Nonetheless, it remains to be investigated how Lm survives within placental and fetal tissue once it is released from neutrophils, by evading defense mechanisms such as those induced by placental Hofbauer cells (Hoo et al., 2024; Yoshida et al., 2025). In summary, our findings reveal for the first time a “Trojan horse” mechanism wherein Lm hijack circulating maternal neutrophils utilizing these immune cells as a survival niche and facilitators of placental and fetal Lm infection.

## Materials and Methods

### Bacterial strains and culture conditions

*Listeria monocytogenes* strains EGDe and EGD constitutively expressing PrfA regulated virulence genes, were grown in BHI medium (BD Biosciences, Franklin Lakes, USA) supplemented with 7 μg/ml chloramphenicol (CM) (VWR, Darmstadt, Germany) at 37 °C and 180 rpm to an optical density (OD_600 nm_) of 0.05-0.5 before included in experiments.

### CFSE staining

CellTrace CFSE Cell Proliferation Kit (C34554) (Invitrogen, Waltham, USA) was used for Lm staining. Lm were grown to an optical density of 0.05-0.5 and 5 x 10^8^, centrifuged, resuspended in 1 ml PBS (Invitrogen, Waltham, USA) stained with 5 µM CFSE (Invitrogen) and incubated for 30 min in the dark at 37 °C. 100 % purity of PBS-washed Lm was checked using FACS analysis.

### Trophoblast cell lines and cell culture

HTR-8/SVneo cells termed HTR-8 cells (ATCC CRL-3271) and JEG-3 cells (kindly provided by Udo Jeschke (Department of Obstetrics and Gynecology, University Hospital Augsburg, Augsburg, Germany) were cultured in RPMI 1640 growth medium (Sigma, Kanagawa, Japan) and supplemented with 10 % FCS (Sigma, Kanagawa, Japan), penicillin and streptomycin (both 100 U/ml) at 37 °C in 5 % CO_2_. Prior to Listeria infection, culture medium was changed to RPMI 1640 growth medium supplemented with 1 % FCS (Sigma) without antibiotics.

### Identification of surface markers relevant for leukocyte recruitment

To check for the expression of surface molecules relevant for leukocyte recruitment, 5 x 10^6^ HTR-8 cells or JEG-3 cells were seeded into flasks, grown overnight and infected with Lm (MOI8) for 2 h. Washed and scraped cells were stained for 20 min at RT with antibodies against CD62E (E-Selectin, clone HAE-1, Biolegend, San Diego, USA), CD62P (P-Selectin, clone Psel.KO2.3, Biolegend, San Diego, USA), CD106 (VCAM-1, clone BBIG-V (4B2), R&D, Minneapolis, USA), CD54 (ICAM-1, clone 84H10, Biolegend, San Diego, USA), CD31 (PECAM, clone 9G11, R&D, Minneapolis, USA), CD182 (CXCR2, BD Biosciences, Franklin Lakes, USA), E-Cad (polyclonal, Cell Signaling, Danvers, USA), and c-Met (monoclonal, Abcam, Cambridge, UK), using a final concentration of 5 µg/ml for primary antibodies, and 1:400 dilution for secondary antibodies. Samples were fixed with 2 % PFA (Merck, Darmstadt, Germany) at RT and analyzed by flow cytometry with a Beckman Coulter Gallios flow cytometer and Kaluza Flow analysis software.

### E-Cadherin and c-Met expression on human trophoblast cells

HTR-8/SVneo cells and JEG-3 cells were seeded on gelatin (Life technologies, Carlsbad, UK) coated coverslips and grown ON. Trophoblast cells were then fixed with 2 % PFA (Merck) for 10 min, blocked and permeabilized with 2 % PBS/BSA (Invitrogen) and 0.1 % TritonX (AppliChem, Chicago, USA) for 1 h at RT. After staining the cells with antibodies against E-Cad (polyclonal, Cell Signaling), and c-Met (monoclonal, Abcam), an Alexa488-conjugated donkey anti-rabbit antibody (Invitrogen, Waltham, USA) was applied for 1 h at RT. All antibodies were used in a final concentration of 5 µg/ml. Nuclei were stained for 5 min at RT with DAPI (Invitrogen, Waltham, USA) and samples embedded in ProLong Diamond antifade mounting medium (Invitrogen, Waltham, USA). A Leica SP8X WLL microscope equipped with a HC PL APO 40x /1.30NA oil immersion objective (Leica, Wetzlar, Germany) at the core facility Bioimaging of the Biomedical Center, LMU Munich, was used for image acquisition. Images were analyzed using ImageJ.

### Human blood sample processing

Healthy female volunteers of the same age donated blood for analysis, which was approved by the local ethical committee of Ludwig-Maximilians-Universität München, Munich, Germany (Az. 611-15, 22-0534). Blood cell counts including leukocyte differential counts were obtained using an automated cell counter (ProCyte Dx; IDEXX Laboratories).

### Isolation of human neutrophils and lymphocytes

Isolation of uninfected human neutrophils was conducted via density centrifugation. Therefore, whole blood was applied onto a layer of Polymorphprep (Axis-shield, Dundee, UK) and centrifuged at 500g for 30 min at room temperature (RT). The cell layer containing neutrophils was collected in a Falcon tube, washed with PBS (Invitrogen), and resuspended in HBSS (Merck, Darmstadt, Germany). Lm-infected human blood was isolated using the EasySep Direct Human Neutrophil Isolation Kit (Stem Cell, Vancouver, Canada) and negative selection of neutrophils performed according to the manufacturer’s instructions.

Isolation of uninfected human lymphocytes was conducted via density centrifugation. After 1:1 dilution of whole blood with PBS (Invitrogen) the samples were applied on a layer of Lymphoprep (Axis-shield, Dundee, UK) and centrifuged at 800g for 20 min at room temperature (RT). The layer containing lymphocytes was collected in a Falcon tube, washed with PBS (Invitrogen) and resuspended in HBSS (Merck).

### Shuttle screening

Two alternative approaches were performed to screen for a potential blood cell shuttle of Lm to the placenta. In a first approach, whole blood from human female donors was infected for 1 h at 37 °C and 120 rpm with CFSE (Invitrogen) + Listeria (MOI1). After gentamicin (100 µg/ml) (Sigma, Kanagawa, Japan) treatment, samples were washed and stained with antibodies against CD14 (Monocytes, clone HCD14, Biolegend, San Diego, USA), CD41 (Platelets, clone HIP8, Biolegend, San Diego, USA), CD56 (Natural killer cells, clone HCD56, Biolegend, San Diego, USA) and CD66b (Neutrophils, clone G10F5, Biolegend, San Diego, USA) using a final concentration of 5 µg/ml. For FACS analysis, samples were fixed with 1.5 % PFA (Merck) and erythrocytes lysed using BD FACS Lysing Solution (BD Biosciences, Franklin Lakes, USA). Intracellular bacterial burden was analyzed with a Beckman Coulter Gallios flow cytometer and Kaluza Flow analysis software. In an alternative approach, neutrophils and PBMCs of human female donors were isolated via respective density centrifugation. Isolated cells were infected with Listeria (MOI8) and treated and stained according to the first approach described above. Viability of intracellular Listeria was maintained via plating of the cell dilutions on agar dishes supplemented with CM (7 µg/ml) (VWR). Counting of CFUs was performed on the next day.

### Heat inactivation of human serum

For investigation of factors in human serum responsible for Lm internalization of neutrophils, whole blood from female donors was taken and used for density centrifugation to isolate neutrophils and to extract serum via centrifugation for 10 min at 2000g. Serum samples were treated for 30 min at 56 °C and 450 rpm for heat inactivation (Drevets and Campbell, 1991). CFSE (Invitrogen) stained Listeria were used for infection of 30 min of isolated neutrophils (MOI8) that were opsonized with the heat inactivated serum, untreated serum and HBSS (Merck) as a control prior to infection. After gentamicin (100 µg/ml) (Sigma) treatment, samples were washed, fixed with 1.5 % PFA (Merck) and intracellular bacterial burden quantified with a CytoFLEX S cytometer and FlowJo analysis software.

### Inactivation of C3 with Cobra Venom Factor

Prior to Listeria opsonization, human serum was de-complemented using 12 µg of cobra venom factor (CVF) (Quidel, San Diego, USA) per 500 µl of extracted serum and incubated for 1 h at 37 °C (Haihua et al., 2018). CFSE (Invitrogen) stained Listeria was used to infect isolated neutrophils (MOI8) opsonized with CVF (Quidel) treated serum, untreated serum, and HBSS (Merck) as a control. After gentamicin (100 µg/ml) (Sigma) treatment, samples were washed, fixed with 1.5 % PFA (Merck) and intracellular bacterial burden quantified with a CytoFLEX S cytometer and FlowJo analysis software.

### Blocking of c-Met receptor and Fc-gamma receptor

c-Met receptor and FcyR blocking on human neutrophils were conducted via respective antibody treatment of isolated neutrophils for 30 min at 37 °C and 120 rpm, respectively. PBS (Invitrogen) was used as control as well as a blocking c-Met ab (polyclonal, Thermo Scientific, Waltham, USA), FcγR blocking ab (Biolgened, San Diego, USA) and blocking c-Met ab (polyclonal, Thermo Scientific) in combination with FcγR blocking ab (Biolegend) at a final concentration of 5 µg/ml. Using a MOI8 CFSE (Invitrogen) stained Listeria were applied to infect isolated pre-treated neutrophils for 30 min. Prior to infection cells were opsonized with untreated serum for 30 min. After gentamicin (100 µg/ml) (Sigma) treatment, samples were washed, fixed with 1.5 % PFA (Merck) and intracellular bacterial burden quantified with a CytoFLEX S cytometer and FlowJo analysis software.

### Shuttle infection of human trophoblasts

1 x 10^6^ HTR8/ SVneo cells or JEG3 cells per well were seeded into gelatin (Life technologies) coated 6-well plates and stimulated with the TLR1/2 agonist PAM3CSK4 (InvivoGen, San Diego, USA) or PBS (Invitrogen) as control for 6 h on the next day. After infection of whole blood from human female donors with CFSE (Invitrogen) stained Listeria (MOI8) for 1 h at 37 °C and 120 rpm, gentamicin (100 µg/ml) (Sigma) was added for 30 min at 37 °C and 120 rpm. Samples were washed with PBS (Invitrogen) and infected neutrophils isolated using the EasySep Direct Human Neutrophil Isolation Kit (Stemcell, Vancouver, Canada). EDTA (1mM final concentration) was added to the blood sample and negative selection of neutrophils was performed according to the manufacturer’s instructions. One half of infected neutrophils was lysed in deionized water to have a free, but comparable amount of Lm to the other half of isolated infected cells that was resuspended in HBSS (Merck) and thus kept functional. Platelet rich plasma was generated from 5 ml of Lm infected blood prior to gentamicin (Sigma) treatment. Using a MOI 6, free Lm that was exposed to neutrophils but lysed with water to release Lm, Lm infected neutrophils, Lm infected platelets and untreated Listeria were applied for 1 h at 37 °C on TLR1/2 agonist-stimulated or unstimulated trophoblast cells, respectively. After gentamicin (100 µg/ml) (Sigma) treatment, wells were washed, trophoblasts were scraped in PBS (Invitrogen) and one part of the cell dilutions plated on agar dishes were lysed in water and checked for Lm burden by counting CFUs on the next day. The rest of the cell dilutions were fixed with 1.5 % PFA, stained with antibodies (5 µg/ml) against CD54 (ICAM-1, clone 84H10, Biolegend) and CD66b (Neutrophils, clone G10F5, Biolegend) for 20 min at RT and intracellular bacterial burden was quantified with a BD LSR Fortessa flow cytometer and FlowJo analysis Software.

### Mice handling

KIE16P mice were provided by Marc Lecuit, Pasteur Institute, Paris, France (Disson *et al*., 2008). C57BL/6 (Nomenclature C57BL/6NCrl) mice from Charles River were included in experiments as WT controls. For the intravital microscopy placenta model, KIE16P mice were crossed with Ly-6A (Sca1) GFP transgenic mice (Ma et al., 2002). Animals were included in experiments at an age of 7-25 weeks and murine fetuses between embryonic days 13.5-15.5. All protocols were reviewed and approved by the government of Oberbayern, Germany, AZ ROB 55.2-1-54-2531-122/12, and -229/15, ROB-55.2-2532.Vet_02-18-26 and 02-23-144, and AZ 50-8791-14.835.2259.

### Intravital microscopy placenta model

Multiphoton intravital microscopy (MPM-IVM) was used to study the immunological environment of the murine placenta *in vivo*. For imaging maternal and fetal blood circulation, a genetic mouse model was created by crossing KIE16P females with Sca1-eGFP mice (provided by C. Robin, Hubrecht Institute, NL), labeling the fetal microvasculature and blood cells with green fluorescent protein (GFP). Maternal blood circulation was visualized by intravenous injection of fluorescent dye (Blue dextran or TRITC, 200 µl at 1:100 dilution from Thermo Fisher, Waltham, USA). To track Lm in placental circulation, we used strains expressing GFP (EGDe_GFP) or Tomato fluorescent protein (EGDe_tomato). Anesthetized mice were intubated, and catheters were placed in the carotid artery (for bacterial and substance infusion) and the tail vein (for dye infusion). The placenta was surgically exposed and positioned on a custom stage for imaging, keeping the embryo connected and covered with warm ultrasound gel to prevent tissue drying. A vacuum suction ring with a cover slip was used to stabilize the tissue (Rodriguez-Tirado et al., 2016), allowing imaging through the cover glass positioned between the placenta and yolk sac. Lm was infused via the carotid artery catheter to capture initial interactions upon arrival at maternal vessels. MPM-IVM images were acquired using the LaVision BioTech TriMScope II multiphoton microscope with a 16x objective (NA 0.8, Nikon) and 3D time-lapse over one to three hours. Images were recorded with a 2 µm z-interval across a 512 x 512 µm field of view. The system was equipped with a Ti;Sa Chameleon Unltra II Laser (Coherent), using laser excitations at 800 nm (Ti;SA) and 1100 nm (OPO) Ti;Sa. Data was captured using ImSpector Pro (LaVision) software, version 275.

### Infection experiments of mouse peripheral blood *in vivo*

Pregnant and non-pregnant anesthetized C57BL/6 mice were injected i.v. with 4 x 10^6^ Listeria of CFSE-labelled EGDe-GFP strain (Lm-infected) or not injected with Lm as mock controls. Time-dependent infection experiments ranged from 10 min to 60 min. After the corresponding time point of infection, blood was harvested by retro-orbital bleeding and mice sacrificed. Immediately after collection, the blood was treated with gentamicin (100 µg/ml) (Sigma) for 30 minutes, washed and then the samples split for FACS analysis and CFU plating. For FACS analysis, blood samples were stained with viability dye (Live/Dead Fixable Blue, Invitrogen) to check for dead cells and then erythrocytes lysed and fixed (BD FACS). Samples were stained with antibodies against CD45-PE-Dazzle 594 (Leukocytes, clone 30-F11, Biolegend, San Diego, USA), CD19-PE (B-cells, clone 6D5, Biolegend, San Diego, USA), CD3e-PE (T-cells, clone 145-2C11, Biolegend, San Diego, USA), Ly6G-PE-Cy7 (Neutrophils, clone 1A8, Biolegend, San Diego, USA), CD11b-BV650 (Myeloid cells, clone M1/70, Biolegend, San Diego, USA), Ly6C-APC-Cy7 (Monocytes, clone HK1.4, Biolegend, San Diego, USA), at a final concentration of 2 µg/ml. Cells were acquired on a CytoFlex S flow cytometer (Beckmann Coulter) and analyzed using FlowJo software. Neutrophils were defined as CD45+ CD3-CD19-Ly6G+. Monocytes were defined as CD45+ CD3-CD19-Ly6G- CD11b+ Ly6C+. CFSE-positive Listeria gates were defined for each immune cell subset based on mock control fluorescence and Listeria uptake was shown relative to mock controls.

For CFU plating, the rest of the blood sample was resuspended with 10x volume of deionized water, incubated for 5 minutes at room temperature, centrifuged at 4000 rpm for 5 minutes and then resuspended in PBS for CFU platings with serial dilutions onto BHI agar. The efficiency of water lysis of immune cells was confirmed with FACS analysis of CD45 stained-lysed samples, showing no intact leukocytes. To quantify the intracellular bacterial burden in peripheral blood, colonies grown on BHI agar plates were counted 24 hours after plating and expressed as CFU per ml of blood.

Due to the low-dose infection and corresponding lower number of infected cells in physiological *in vivo* infections, CFU plating of whole blood from pregnant mice were compared with non-pregnant mice and microscopy of pregnant mice leukocytes were done to observe infected and viable Lm-immune cells populations.

### Confocal microscopy of Listeria in immune cells of mouse peripheral blood

Pregnant or non-pregnant mice were infected as above for the FACS experiments with higher dose of 4 x 10^8^ to ensure sufficient number of infected cells for visualization. Peripheral blood was harvested and treated with gentamicin (100 µg/ml, 30 minutes). Following erythrocyte lysis and fixation, leukocyte populations were seeded onto 8-well removable glass dishes (Ibidi) pre-coated with 0.01% poly-L-lysine (Sigma Aldrich). After 2 hours, adhered cells were fixed with 4% PFA (Merck) for 10 minutes at room temperature and then permeabilized with 0.1% Triton-X-100 in 2% BSA-PBS for 10 minutes, followed by blocking in 2% BSA-PBS for 30 minutes. Following washes, the cells were then stained overnight with a combination of the following primary antibodies against Ly6G (Neutrophil membrane marker, clone EPR22 909-135, Abcam), Ly6C (Monocyte membrane marker, clone ER-MP20, Abcam), LAMP-1 (Lysosomal membrane marker, clone H4A3, Invitrogen), Galectin-3 (phagosomal rupture marker, polyclonal, Biotechne), at a final concentration of 2-5 µg/ml. The cells were then stained the next day with the corresponding fluorescent dye labelled secondary antibodies for 1 hour: donkey anti-Alexa 546, donkey anti-mouse Alexa 594, and donkey anti-rat Alexa 647 or donkey anti-goat Alexa 647. DAPI (Invitrogen) was used to stain the nuclei afterwards for 5 minutes, washed, and samples were embedded on glass slides with VectaShield Antifade mounting medium (Vector Laboratories). A Leica SP8X WLL microscope equipped with a 63x 1.4NA oil immersion objective (Leica, Wetzlar, Germany) at the core facility Bioimaging of the Biomedical Center, LMU Munich, was used for image acquisition. Five-color images were acquired with sequential imaging to avoid bleed-through and the image pixel size was 70 nm. Z-stack projections were recorded for all samples and images were analyzed using ImageJ software. All images intended to be compared, linearly contrast stretched to display relevant features and processed identically to ensure there was no bias. Quantification of Galectin-3 punctae per cell was performed on maximum intensity projections of z-stack of infected neutrophils and monocytes identified by Listeria+Ly6G+ channels and Listeria+Ly6C+ channels, respectively. Additional quality control was done by checking the DAPI channel of each image. The Galectin-3 channel was converted to binary images after background subtraction and strict thresholding. Particles were analyzed based on size and circularity cutoffs. Cells with more than 2 Gal-3 punctae detected were considered to be Gal-3 positive. Thereafter, Gal-3 positive cells were checked for colocalization with CFSE-Listeria signal, which then indicated cytosolic escape of Lm. Only infected cells from the same experiment with three mice were compared to ensure uniform staining and imaging conditions.

### Infection experiments of mouse maternal-fetal invasion *in vivo*

Pregnant KIE16P or C57BL/6 mice were infected i.v. with 4 x 10^6^ Listeria of the EGDe strain in 100 µl PBS (Invitrogen) after neutrophil depletion, blocking neutrophil adhesion or without pretreatment. Depletion of neutrophils in mice was performed by i.v injection of anti Ly6G monoclonal antibody (clone 1A8, 100 µg per injection, Biolegend, San Diego, USA) 24 h and 4 h before the experiment started. Successful depletion of the neutrophil population was confirmed via FACS analysis of peripheral blood samples. For blocking of neutrophil adhesion in the placenta *in vivo*, antibodies against CD11b/CD18 (MAC-1, clone TIB128, 100 µg, LGC, Teddington, UK) and CD11a (LFA-1, clone TIB217, 30 µg, LGC, Teddington, UK) were admistered i.p. 2 h before Lm injection. Via infection of KIE16P mice with an EGDe strain lacking Internalin A, a functional role of the humanized E-Cadherin receptor in the murine placenta was investigated. Eight hours after infection with Lm, mice were sacrificed, maternal liver (control organ), placentas and fetuses were removed, washed and homogenized. After gentamicin (100 µg/ml) (Sigma) treatment for 30 min samples were washed with PBS (Invitrogen) and incubated with deionized water for 10 min for cell lysis of the respective organ. To quantify the bacterial burden in the placental fetal unit (PFU) and evaluate susceptibility to Lm during pregnancy, dilutions of smashed organs were cultured on BHI agar (BD Biosciences) dishes and CFUs were counted the days after the experiment.

### Transmission electron microscopy

Samples were fixed with 2.5% glutaraldehyde in 0.1M sodium cacodylate buffer, pH 7.4 (Electron Microscopy Sciences, USA) for 24 h at minimum. Thereafter glutaraldehyde was removed and samples were washed three times with 0.1M sodium cacodylate buffer, pH 7.4. Postfixation and prestaining was done for 45 to 60 min with 1% osmium tetroxide (Electron Microscopy Sciences, USA), Samples were washed three times with ddH_2_O and dehydrated with an ascending ethanol series (15min with 30%, 50%, 70%, and 90% respectively and two times 100% over 10min). Subsequently, samples were embedded in Epon (Serva Electrophoresis GmbH). 60-70 nm thick ultrathin sections were cut at the Reichard-Jung Ultracut E microtome (Darmstadt, Germany). Ultrathin sections were collected on formvar coated copper grids (Plano, Germany) and automatically stained with UranyLess EM Stain (Electron Microscopy Sciences) and 3% lead citrate (Leica, Wetzlar, Germany) using the contrasting system Leica EM AC20 (Leica, Wetzlar, Germany). Imaging was carried out using the JEOL -1200 EXII transmission electron microscope (JEOL, Akishima, Tokyo) at 80 kV. Images were taken using a digital camera (KeenViewII; Olympus, Germany) and processed with the iTEM software package (anlySISFive; Olympus, Germany).

### Tissue clearing for infected placenta imaging and analysis

To investigate Lm – neutrophil colocalization in the placenta of Lm infected pregnant mice, we applied a tissue clearing approach of control and infected placentas. For this, we infected pregnant KIE16P mice (E12.5-15.5) via intraarterial injection of Lm constitutively expressing a GFP (EGDe_GFP) into the carotid artery. After 60 min incubation, mice were sacrificed and placentas surgically removed, washed, and prepared for tissue clearing. First, whole placentas were dehydrated with stepwise methanol addition in H₂O (50% for 1 h, 80% for 1 h, and 100% for 2 h each). They were then incubated overnight at 4°C in 66% dichloromethane/34% methanol. After this, samples were washed twice in 100% methanol for 1 h each. Overnight bleaching was performed at 4°C using 5% H₂O₂ (1 volume 30% H₂O₂ to 5 volumes methanol). Following bleaching, the placentas were gradually rehydrated in 80%, 50%, and 20% methanol in H₂O for 1 h each step, followed by two washes in PBS. Permeabilization was then conducted in PBS with 0.2% Triton X-100 for 2 h, after which the placentas were incubated overnight at 37°C in PBS with 0.2% Triton X-100, 10% DMSO, and 0.3 M glycine. Blocking was performed in PBS with 0.2% Triton X-100, 10% DMSO, and 6% goat serum for 1 day. After blocking, the placentas were incubated with Ly-6G PE antibody (Invitrogen, 1A8-Ly6g, Catalog #12-9668-82, dilution 1:1000) at 37°C in PBS with 0.2% Tween-20 and 10 mg/L heparin (PTwH) for 5 days. The samples were then washed three times for 3 h each at 37°C in PTwH. Next, they were incubated with GFP-Booster Alexa Fluor 647 (ChromoTek, Catalog # gb2AF647, dilution 1:1000) and Goat anti-Rat IgG (H+L) Cross-Adsorbed Secondary Antibody, Alexa Fluor 568 (Invitrogen, Catalog # A-11077, dilution 1:1000) at 37°C in PTwH for 5 days. The samples were then washed three times for 3 h each at 37°C in PTwH. Following antibody incubation, the placentas were gradually dehydrated using a tetrahydrofuran (THF) gradient in distilled water (50%, 70%, 90%, and 100% for 2 h each, with an additional overnight incubation in 100% THF). They were subsequently incubated for 1 h in dichloromethane (DCM, Sigma, 270997) and finally cleared in BABB (1 part benzyl alcohol to 2 parts benzyl benzoate, Sigma, 24122 and W213802) until optical transparency was achieved.

### Statistical analysis

Data in this study were analyzed and edited using GraphPad Prism 10 software. All data were depicted as either mean±SEM, cumulative frequencies, median or representative images and plots. Depending on the number of groups that were compared, respective statistical tests were applied. Unpaired or paired student’s t-test were used to compare two groups. In case of more than two groups, a 1-way or 2-way analysis of variance (ANOVA) with either Dunnett’s (comparison of experimental groups against control) or Tukey’s (comparison of all groups with each other) were conducted. Statistical significance was assessed as follows: *p <0.05; **p <0.01; ***p<0.001; ****p<0.0001. Figures were assembled for publication using Inkscape (San Jose, CA, USA). Animations were created with BioRender.com.

## Data availability statement

## Acknowledgements

We thank Dorothee Gössel and Susanne Bierschenk for their excellent technical assistance. We also thank Werner Goebel and Wolfgang Eisenreich (both LMU Munich) for their valuable comments and suggestions during this project. We are indebted to the multiphoton imaging core facility at the Walter Brendel Centre of experimental Medicine, LMU Munich, Germany, the core facility flow cytometry as well as the core facility bioimaging at the Biomedical Center, LMU Munich, Planegg-Martinsried, Germany. We thank Catherine Robin, Hubrecht Institute, Utrecht, NL, for kindly providing Sca1-eGFP mice. We would also like to acknowledge Lis de Weerd and Christian Haass (DZNE, Munich) for providing us with the Galectin-3 antibody.

## Conflict of interest statement

All authors have declared that no conflict of interest exists.

## Funding

Work was supported by the German Research Foundation (DFG) collaborative research grant SFB914, project Z01 (to H.I.A. and S.M.), project B05 (to R.H.) and projects B01 and Z03 (to M.S.), and TRR359 grant #491676693, project B02 (to R.I. and M.S.).

## Notes

### Competing Interest Statement

The authors have declared no competing interest.

