## Supplemental material for "Neutrophils are critical for placental and fetal infection with the human pathogen *Listeria monocytogenes*"

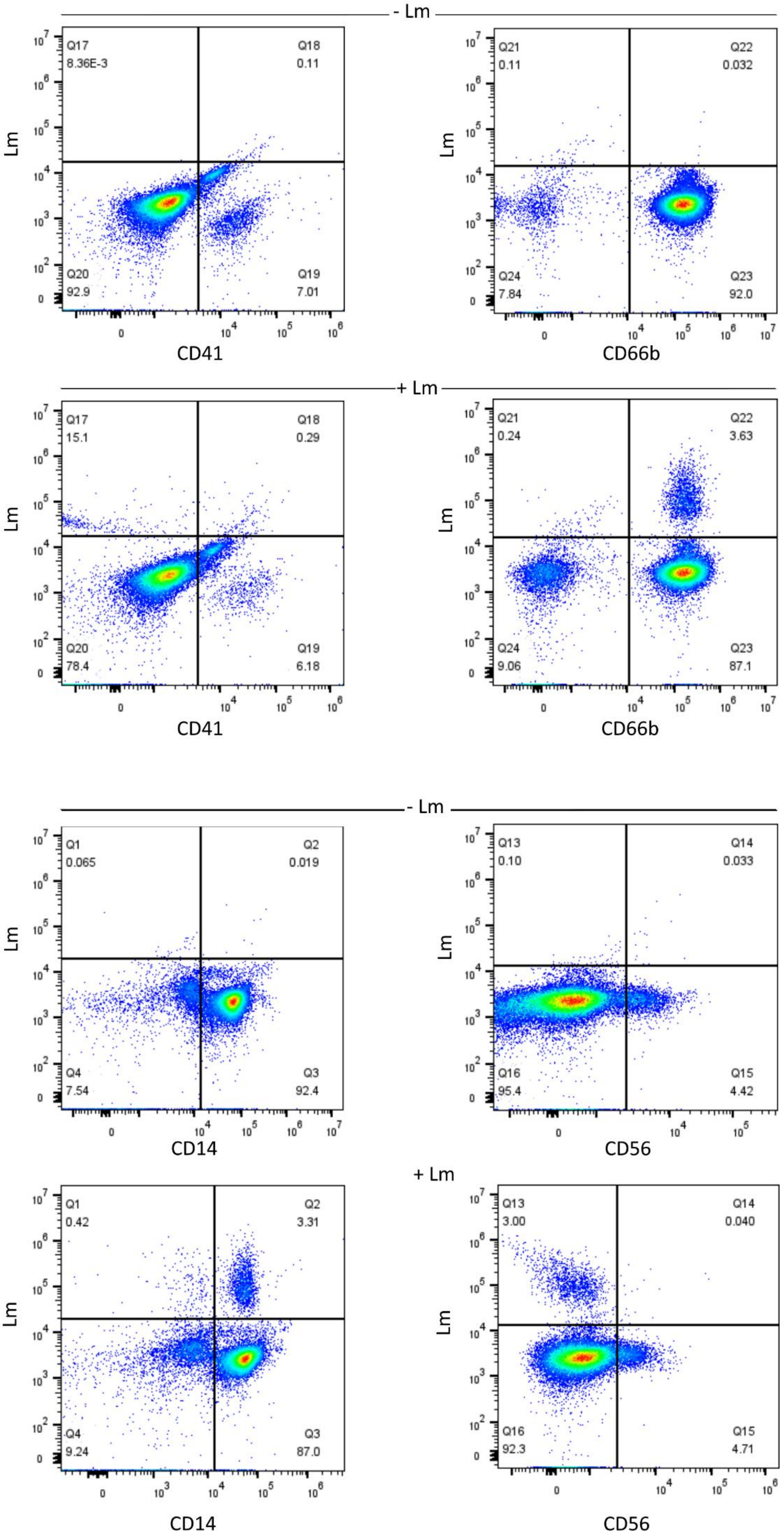

- Lm
+ Lm
Lm
CD41
Q17 8.36E-3
Q18 0.11
Q20 92.9
Q19 7.01
CD66b
Q21 0.11
Q22 0.032
Q24 7.84
Q23 92.0
Q17 15.1
Q18 0.29
Q20 78.4
Q19 6.18
Q21 0.24
Q22 3.63
Q24 9.06
Q23 87.1
CD14
Q1 0.065
Q2 0.019
Q4 7.54
Q3 92.4
CD56
Q13 0.10
Q14 0.033
Q16 95.4
Q15 4.42
Q1 0.42
Q2 3.31
Q4 9.24
Q3 87.0
Q13 3.00
Q14 0.040
Q16 92.3
Q15 4.71


1

**Supplemental Figure 1. FACS analysis of Lm uptake into human peripheral blood cells.** Gating strategy to identify platelets (CD41), neutrophils (CD66b), NKs (CD56) andmonocytes (CD14) from isolated human peripheral blood cells, before and after infection with Lm, as in Fig. 2B. Cell populations that are positive for Lm marker and the respective marker of the cell population, as indicated in the top right quadrants, were included in the analyses. Representative FACS plots are shown from n=4 independent experiments.

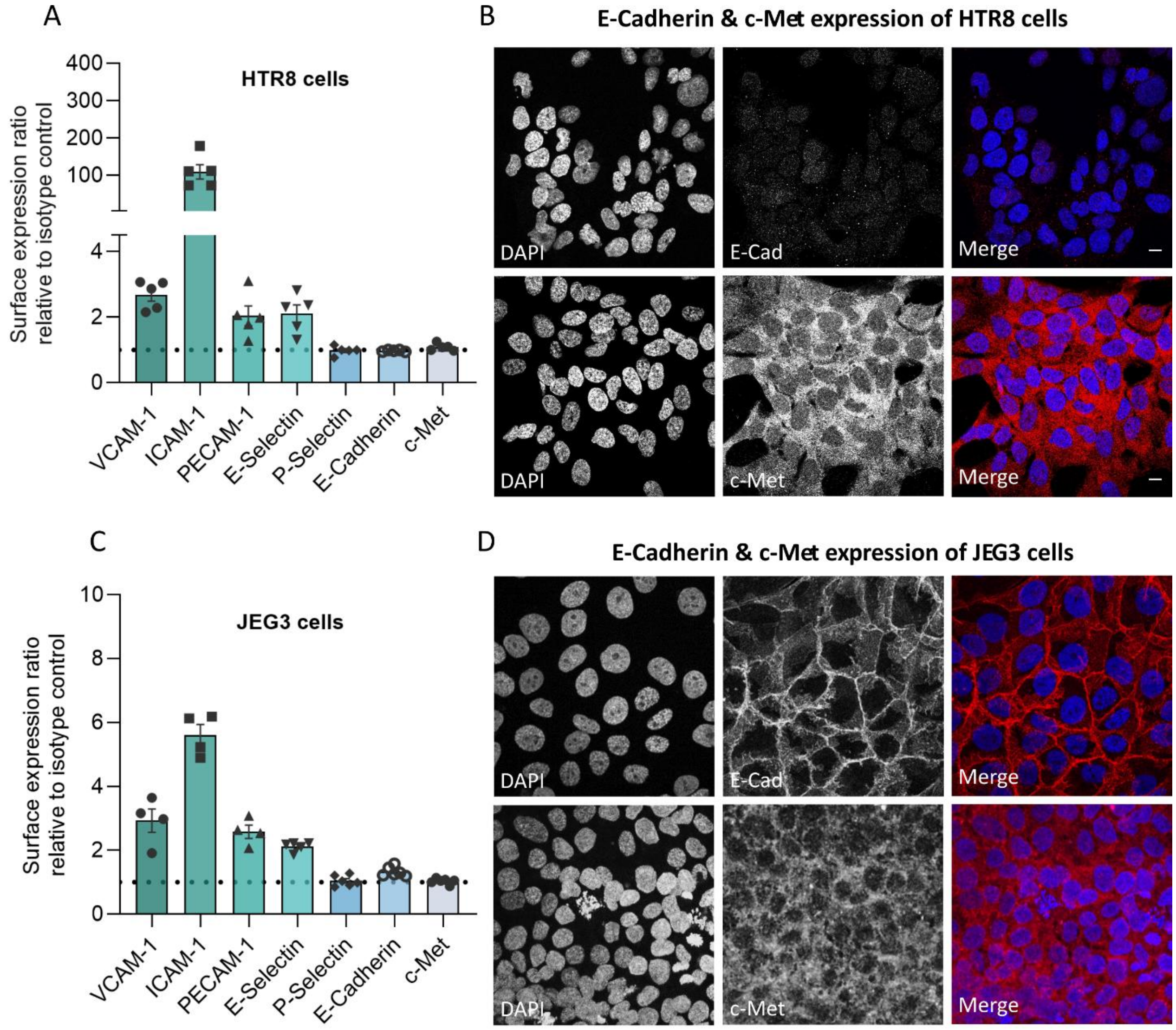


**Supplemental Figure 2. Expression of rolling and adhesion relevant molecules on trophoblast cell lines, HTR8 and JEG3.** Expression of rolling and adhesion relevant molecules as well as expression of E-Cad and c-Met serving as ligands for Lm. Data is shown relative to isotype control on HTR8 (A) and JEG3 (C) trophoblast cells (mean±SEM, n=4-5 independent experiments, 1-way ANOVA, Tukey's multiple comparison). Normalized baseline is indicated with dotted lines in the graphs. Representative confocal images of E-Cad and c-Met expression in HTR8 (B) and JEG3 (D) trophoblast cells are shown (n=3 independent experiments, scale bars = 10 µm).

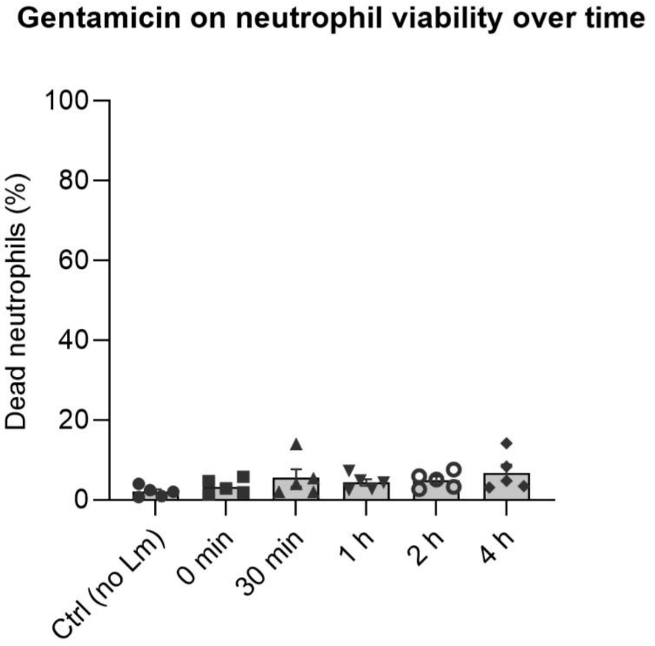


**Supplemental Figure 3. Viability of neutrophils upon Lm infection with gentamicin treatment.** Viability of isolated human neutrophils was assessed during gentamicin treatment (mean±SEM, n=5 independent experiments, no significant changes were observed, 1-way ANOVA, Tukey's multiple comparison).

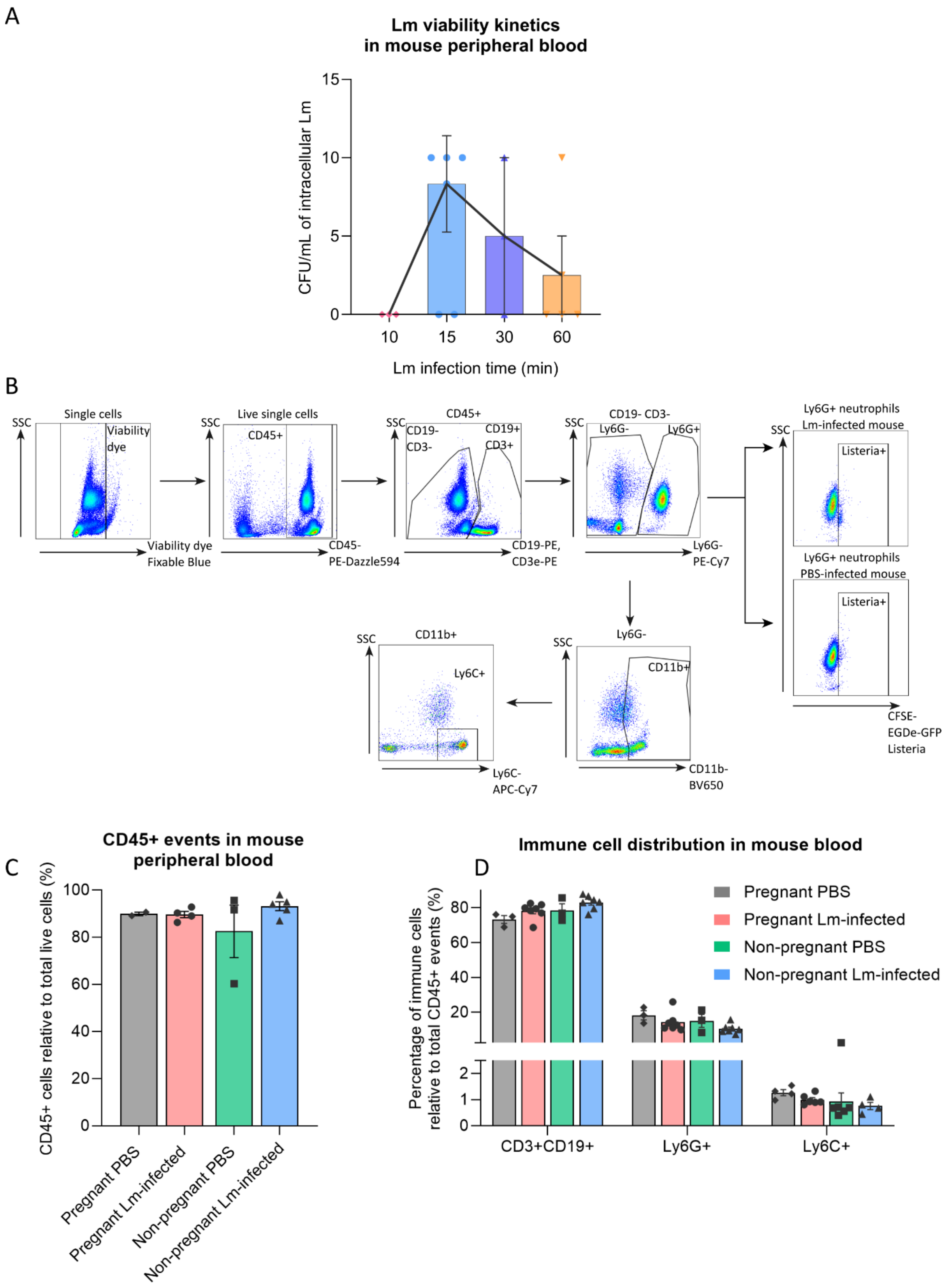
A
Lm viability kinetics
in mouse peripheral blood
CFU/mL of intracellular Lm
15
10
5
0
10
15
30
60
Lm infection time (min)
B
Single cells
SSC
Viability dye
Viability dye
Fixable Blue
Live single cells
CD45+
CD45-
PE-Dazzle594
CD45+
CD19-
CD3-
CD19+
CD3+
CD19-PE,
CD3e-PE
CD19- CD3-
Ly6G-
Ly6G+
Ly6G-
PE-Cy7
Ly6G+ neutrophils
Lm-infected mouse
Listeria+
Ly6G+ neutrophils
PBS-infected mouse
Listeria+
CFSE-
EGDe-GFP
Listeria
Ly6G-
CD11b+
CD11b-
BV650
CD11b+
Ly6C+
Ly6C-
APC-Cy7
C
CD45+ events in mouse
peripheral blood
CD45+ cells relative to total live cells (%)
100
80
60
40
20
0
Pregnant PBS
Pregnant Lm-infected
Non-pregnant PBS
Non-pregnant Lm-infected
D
Immune cell distribution in mouse blood
Percentage of immune cells
relative to total CD45+ events (%)
80
60
40
20
2
1
0
CD3+CD19+
Ly6G+
Ly6C+
Pregnant PBS
Pregnant Lm-infected
Non-pregnant PBS
Non-pregnant Lm-infected

**Supplemental Figure 4. *In vivo* Lm infection experimental setup and FACS analysis of Lm uptake in mouse peripheral blood cells.** (A) CFU counts analysis of intracellular Lm in gentamicin-treated and lysed peripheral blood at different time points after infection. Data is represented as CFU per ml of blood. Black line connects the means of the various groups and shows the highest intracellular Lm viability at 15 min of Lm infection (mean ± SEM, n=2-5 mice per condition) (B) Representative dot plots of the gating strategy used for flow cytometry of Listeria association with the various immune cells of mouse peripheral blood. Markers to identify leukocytes (CD45), B-cells (CD19), T-cells (CD3), neutrophils (Ly6G), myeloid cells (CD11b), and monocytes (Ly6C) were used to define immune cell subsets. Infected cells are further identified by gating on the mock-infected mice to detect the CFSE-Lm-positive population under each immune cell gate and then applied to the infected samples. Dot plots are representative of 7 individual experiments. (C) CD45+ leukocytes are plotted relative to the total live single cells acquired per mouse sample, showing comparable events across groups (mean ± SEM, n=3-7 mice per condition, 2-way ANOVA, Tukey's multiple comparison). (D) Distribution of various immune cell subsets across mouse groups are plotted as percentage of each immune cell to total CD45+ events, to ensure comparable leukocyte distribution. This shows no inherent differences in proportion of immune cells across mice ± infection (mean ± SEM, n=3-7 mice per condition, 2-way ANOVA, Tukey's multiple comparison).

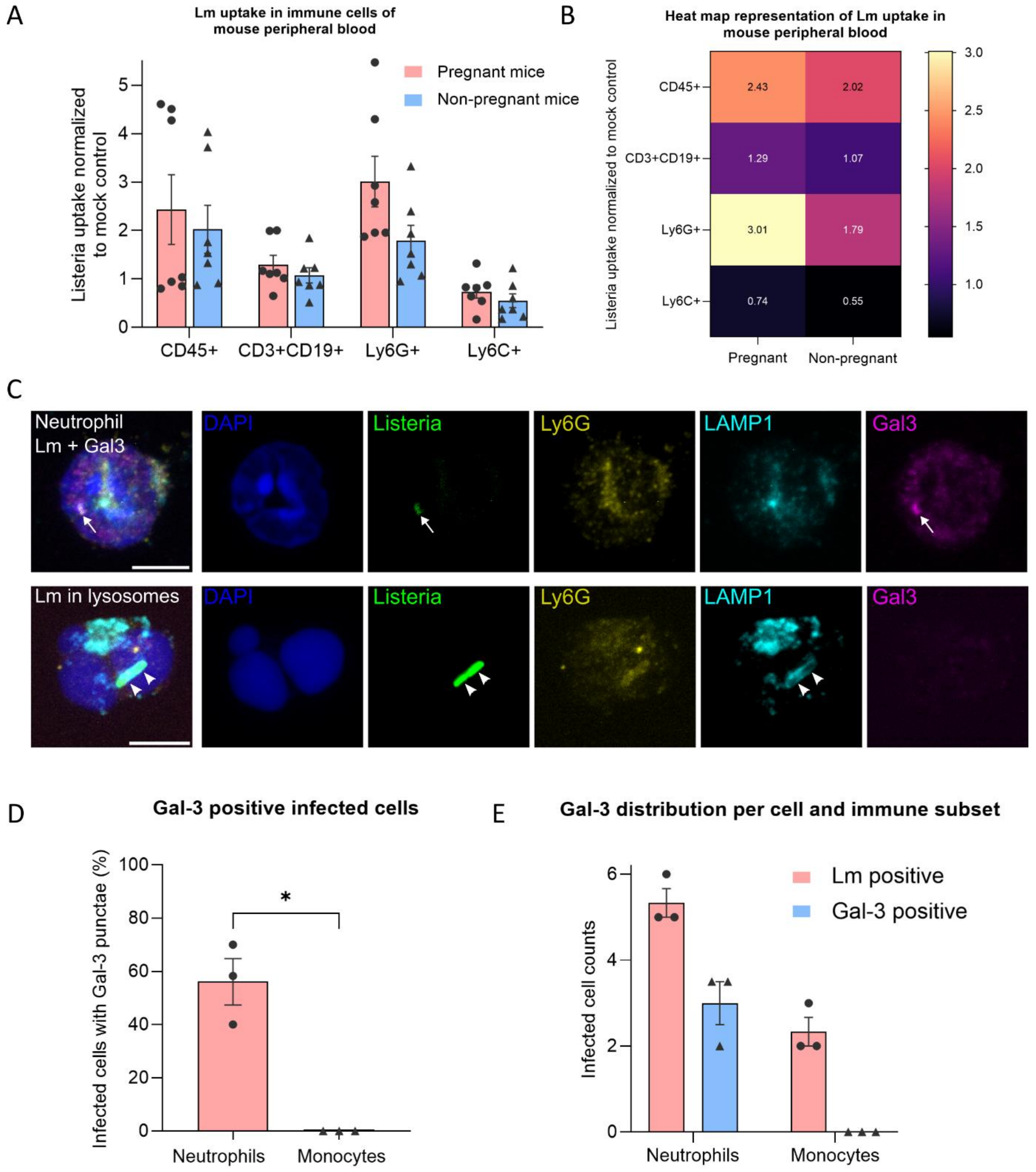


**Supplemental Figure 5. Lm infection *in vivo* shows neutrophils in peripheral blood to act as an intravascular shuttle for Listeria.** (A) FACS analysis of *in vivo* Lm infection of various immune cell subsets in peripheral blood of pregnant and non-pregnant mice. Data are shown for CD45+ leukocytes, CD3+CD19+ lymphocytes, Ly6G+ neutrophils and Ly6C+ monocytes, grouped based on pregnancy status. CFSE-positive Lm uptake in immune cells of infected mice is normalized to corresponding background signal in mock-infected controls (mean ± SEM, n=7 mice per condition, 2-way ANOVA, Tukey's multiple comparison). (B) Heatmap representation of the FACS analysis of Lm uptake shown in (A), highlighting maximal uptake in neutrophils of pregnant mice. Heatmap legend is shown on the right and mean values are indicated on each block (mean, n=7 mice per condition, 2-way ANOVA, Tukey's multiple comparison). (C) Confocal microscopy images showing maximum intensity projections of Lm-infected pregnant mice leukocytes. Top panel shows an example image of infected pregnant mouse neutrophil with Lm showing cytosolic escape (similar to

Figure 5C). Neutrophil is identifiable by nuclear DNA staining (DAPI shown in blue) and neutrophil-specific membrane marker (Ly6G shown in yellow), and internalized Listeria are shown in green. Listeria signal colocalizes with lysosomal escape marker, Galectin-3 (magenta; indicated with white arrows), but not with lysosomal compartments (lysosomal membrane protein LAMP-1 in cyan). Bottom panel labelled ‘Lm in lysosomes’ shows an image of a neutrophil confined with lysosomes indicating no vacuolar escape (white arrowheads point to colocalized Listeria and LAMP1 signal). This is also detectable by the lack of Gal3 expression as compared to the panel above. Scale bars = 5 µm. Images representative of 3-4 independent mice. (D) Percentage of infected cells positive for Galectin-3 punctae count plotted per infected mouse shown for neutrophils and monocytes (mean ± SEM, n=3 mice per condition, unpaired student’s t-test with Welch’s correction). (E) Distribution of infected cell counts for neutrophils and monocytes that are Listeria-positive (red circles) and Listeria-positive with Gal-3 punctae (blue triangles). Individual points plotted show average counts per mouse (mean ± SEM, n=3 mice per condition). *$p < 0.05$.

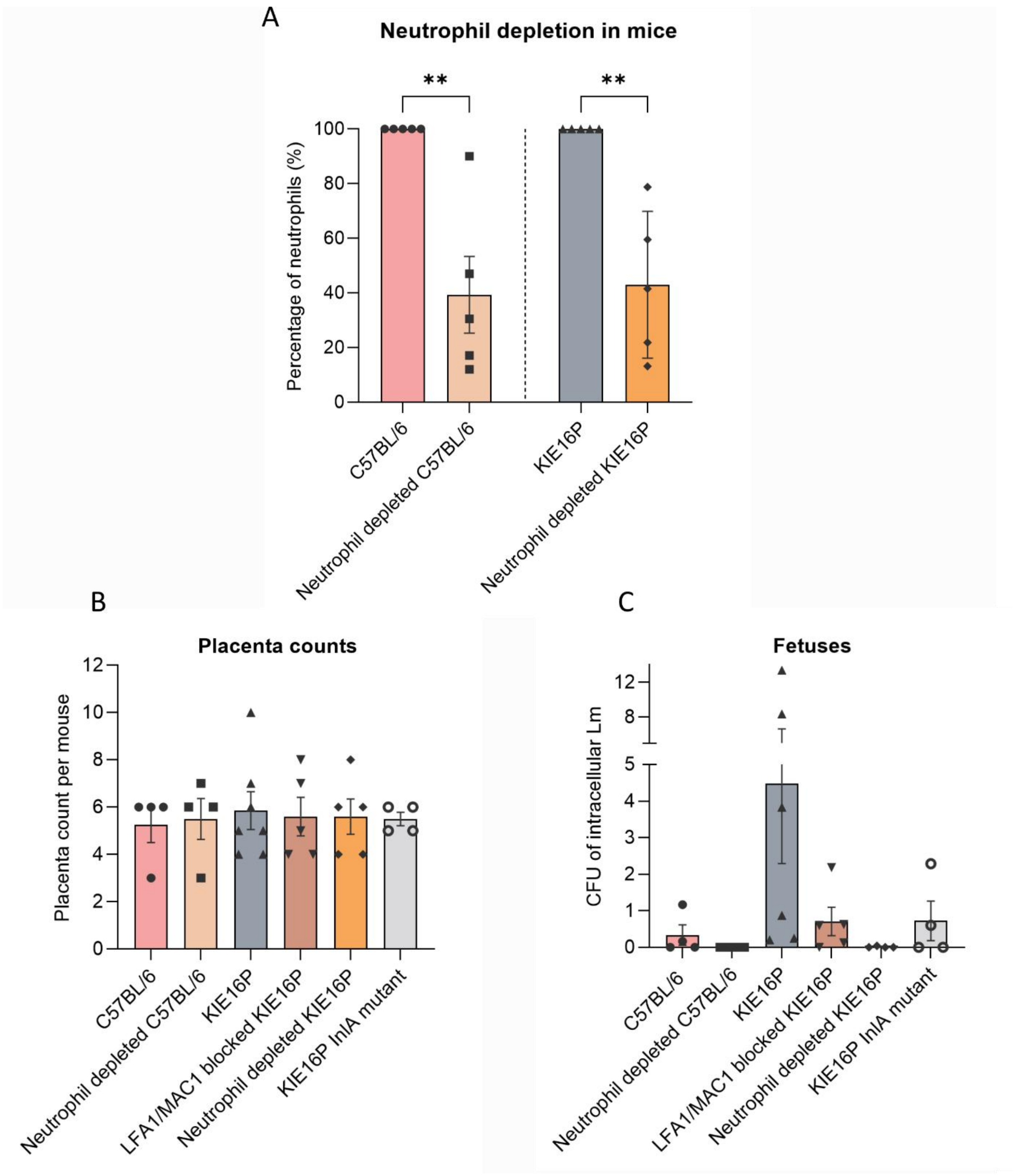


**Supplemental Figure 6. Neutrophil depletion and Lm infection in fetuses and placentas of C57BL/6 and KIE16P mice.** (A) FACS analysis of the percentage of neutrophils before and after depletion in C57BL/6 mice (left side of the graph before the dotted line, n=5 mice) and KIE16P mice (right side of the dotted line, n=5 mice). Mean±SEM for both graphs compared with unpaired student‘s t-test. **p <0.01. (B) Absolute number of placentas per mouse condition plotted showing no major variations across mice (mean±SEM, n=4-5 mice per condition). (C) Listeria load in the fetuses of mice across various conditions (as in Fig. 7) presented in amount of CFUs (mean±SEM, n>4 mice per condition).

**Supplemental Movie 1. Intravital microscopy of *Listeria monocytogenes* (Lm) in the placental microcirculation.**

This video captures the transient presence of Lm in the maternal microvasculature of the murine placenta, visualized using multiphoton intravital microscopy. Maternal blood is labeled with a blue fluorescent dextran dye, delineating the maternal vessels. The left panel shows Lm (red) traveling within the maternal vessels. The right panel presents a 3D reconstruction, highlighting the localization of Lm in the maternal vasculature during the initial minutes post-infection. Clearance of Lm is observed approximately 12 minutes after infection.

**Supplemental Movie 2. Tissue clearing of complete Lm infected mouse placentas.**

3D reconstruction of an intact mouse placenta following *Listeria monocytogenes* (Lm) infection reveals a partial colocalization of Lm and neutrophils in the section view. The background is depicted in green, Lm in blue, and Ly6G-labeled neutrophils in red.
